# Concurrence of Porin Loss and Modular Amplification of β-Lactamase Encoding Genes Drives Carbapenem Resistance in a Cohort of Recurrent *Enterobacterales* Bacteremia

**DOI:** 10.1101/616961

**Authors:** William C. Shropshire, Samuel L. Aitken, Reed Pifer, Jiwoong Kim, Micah M. Bhatti, Xiqi Li, Awdhesh Kalia, Jessica Galloway-Peña, Pranoti Sahasrabhojane, Cesar A. Arias, David E. Greenberg, Blake M. Hanson, Samuel A. Shelburne

## Abstract

**Background:** Carbapenem resistant *Enterobacterales* (CRE) remain urgent antimicrobial resistance threats. Approximately half of CRE clinical isolates lack carbapenem hydrolyzing enzymes and develop carbapenem resistance through alternative mechanisms. The purpose of this study was to elucidate the development of carbapenem resistance mechanisms from clonal, recurrent extended-spectrum β-lactamase positive *Enterobacterales* (ESBL-E) bacteremia isolates in a vulnerable patient population.

**Methods:** This study investigated a historical, retrospective cohort of ESBL-E bacteremia cases in the University of Texas MD Anderson Cancer Center (MDACC) from January 2015 to July 2016. Phylogenetic and comparative genomic analyses were performed to identify clonal, recurrent ESBL-E isolates developing carbapenem resistance. Oxford Nanopore Technology (ONT) long-read and Illumina short-read sequencing data were used to generate consensus assemblies and to identify signatures of mobile genetic element mediated amplification and transposition of antimicrobial resistance genes. Serial passaging experiments were performed on a set of clinical ST131 ESBL-E isolates to recapitulate *in vivo* observations. qPCR and qRT-PCR were used to determine respective copy number and transcript levels of β-lactamase genes.

**Results:** 116 ESBL-E bacteremia cases were identified, 16 of which had documented recurrent infections. Four serial, recurrent isolates displayed a carbapenem resistant phenotype, three without the acquisition of a known carbapenemase. These three isolates had non-carbapenemase-producing CRE (non-CP-CRE) mechanisms driven by IS*26*- and IS*Ecp1*-mediated amplification of respective translocatable units (TU) and transposition units (TPU) harboring both *bla*_OXA-1_ and *bla*_CTX-M_ variants with concomitant outer membrane porin disruption. The TU and TPU structures inserted into the open reading frames of outer membrane porin genes in a subset of non-CP-CRE isolates. Serial passage of an index ST131 ESBL-E isolate under selective carbapenem exposure resulted in chromosomal amplification of modular, TUs harboring β-lactamase genes with concomitant porin inactivation, recapitulating the *in vivo* carbapenem resistance progression. Long-read sequencing of two additional MDACC bacteremia strains identified similar non-CP-CRE mechanisms observed in the serial isolates.

**Conclusions:** Non-CP-CRE *de novo* mechanisms were the primary driver of CRE development in recurrent bacteremia cases within this vulnerable patient population. The incorporation of long-read ONT data into AMR surveillance platforms is critical to identify high-risk CRE isolates that are difficult to identify with low-resolution phenotypic and molecular characterization methods.

## BACKGROUND

Antimicrobial resistance (AMR) is an emerging global health priority and carbapenem resistant *Enterobacterales* (CRE) are among the most serious AMR threats (1). Carbapenem resistance can develop due to the acquisition of enzymes that hydrolyze carbapenems, known as carbapenemases, as well as through changes in outer membrane permeability and/or drug efflux activity, which decrease intracellular carbapenem concentrations (2). While most CRE research has focused on characterizing carbapenemases (2), recent clinical and molecular epidemiology studies indicate approximately 50% of CRE isolates are not carbapenemase carriers, suggesting a substantial proportion of CRE isolates develop carbapenem resistance through alternative mechanisms (3, 4). Molecular characterization of these alternative mechanisms indicate non carbapenemase-producing carbapenem resistant *Enterobacterales* (non-CP-CRE) generally carry extended-spectrum β-lactamases (ESBL) or AmpC-like enzymes with concomitant mutations that alter porin function (2). Further studies reveal that *Enterobacterales* strains carrying outer membrane porin mutations, but lacking ESBL or AmpC-like enzymes, develop *de novo* carbapenem resistance at lower rates during serial passage under increasing carbapenem concentrations (5, 6). Thus, the presence of ESBL or cephalosporinase genes may be a component cause in non-CP-CRE development. Additionally, van Boxtel et al. demonstrated that serial passaging of an ESBL or AmpC-producing isolate in the presence of a carbapenem can result in amplification of plasmid-borne β-lactamase genes (7). Increased expression of the narrow-spectrum TEM β-lactamases has similarly been reported to result in cefepime (8) and piperacillin-tazobactam resistance (9–11), indicating that β-lactamase gene dosage is a factor in increasing resistance to multiple β-lactam chemotherapies. These findings demonstrate expression level and copy number of β-lactamases without known carbapenemase activity have important effects on carbapenem susceptibility in porin deficient backgrounds (6, 7). Nevertheless, there remains a gap in knowledge regarding how these non-CP-CRE mechanisms evolve *in vivo*, as few existing studies have investigated serially collected isolates from large, patient cohorts (12–16).

This gap in knowledge is particularly relevant in carbapenem resistant *Escherichia coli*, where the majority of these isolates are non-CP-CRE (3, 4). Additionally, *Klebsiella pneumoniae* isolates with non-CP-CRE phenotypes have also been identified, albeit less frequently (3, 4). Both non-CP-CR *E. coli* and *K. pneumoniae* can cause severe disease, including bacteremia, which have high mortality rates (17). The detection and treatment of infections with these non-CP-CRE isolates remain challenging relative to carbapenemase producing CRE as phenotypic tests may incorrectly identify these isolates as carbapenem susceptible and definitive therapy options may not be readily evident. This highlights the importance to further characterize non-CP-CRE mechanism development, which may provide insights into new surveillance and/or treatment options. Since there remains a lack of studies fully characterizing non-CP-CRE development, we performed a systematic analysis of non-CP-CRE mechanisms using a large cohort of patients with whole genome sequencing (WGS).

We utilized a cohort of patients with ESBL-E bacteremia from the University of Texas MD Anderson Cancer Center (MDACC). Importantly, analysis of a previous MDACC cohort of non-CP-CRE bacteremia isolates indicated these strains had increased short-read mapping of β-lactamase encoding genes suggestive of β-lactamase encoding gene amplification (18). Nevertheless, the amplification structures and genomic context of these gene amplifications could not be discerned with this short-read sequencing analysis. This is due to inherent limitations with short-read sequencing assemblers, which have difficulties resolving the complex, repetitive mobile genetic elements (MGEs) that carry resistance genes (19). Therefore, we present an analysis utilizing Oxford Nanopore Technologies (ONT) MinION sequencing, a long-read sequencing platform, which overcomes these limitations to elucidate these amplification mechanisms. This study aims to track the *de novo* development of carbapenem resistance mechanisms during recurrent *Enterobacterales* bacteremia infection by specifically focusing on ESBL positive *E. coli* and *K. pneumoniae* strains and characterizing the non-CP-CRE associated β-lactamase amplification mechanisms within our cohort.

## METHODS

### Study design and clinical data abstraction

A retrospective review of patients with ESBL-E bacteremia hospitalized from January 2015 to July 2016 was conducted at MDACC in Houston, Texas. All patients with one or more episodes of ESBL-E bacteremia who were 18 years of age or greater were eligible for inclusion. Blood culture isolates are routinely saved at MDACC and stored at −80°C. Clinical and demographic characteristics were manually extracted from electronic medical records and recorded using REDCap software (Vanderbilt University, Nashville, TN) (20).

### Isolate identification and antimicrobial susceptibility testing

Antibiotic susceptibility testing was performed per routine clinical laboratory practice using an automated system (Vitek2, bioMérieux, Marcy L’Étoile, France) with additional testing performed as needed using individual antibiotic gradient strips (Etest, bioMérieux). ESBL production was assessed per routine laboratory practice on *E. coli*, *K. pneumonia*, and *K. oxytoca* isolates that were resistant to one or more oxyimino-cephalosporins (e.g. cefotaxime, ceftriaxone, or ceftazidime) using either the ESBL Etest (bioMérieux) or the Rapid ESBL Screen kit (ROSCO, Taastrup, Denmark). Carbapenemase production was evaluated in the clinical lab on any *Enterobacterales* isolate resistant to one more of the carbapenems using the Neo-Rapid CARB kit (ROSCO) according to manufacturer’s instructions. Carbapenem resistance (CR) was defined as resistance to either ertapenem or meropenem using CLSI criteria (21). Recurrent *Enterobacterales* bacteremia was defined as identification of the same species in blood culture at any point during the follow-up period following at least one negative blood culture and completion of an antibiotic treatment regimen.

### Illumina short-read sequencing

All available isolates from recurrent bacteremia patients, initially underwent whole genome sequencing (WGS) via Illumina HiSeq as described previously (18). The paired-end short-reads were assessed using the FastQC toolkit (Babraham Institute), and adaptors as well as low-quality reads were trimmed using Trimmomatic v0.33 (22). Genome assembly was performed using SPAdes v3.9.1 (23). Depth of short read mapping to individual genes of interest was quantitated relative to the average read mapping depth for the pubMLST housekeeping gene schema for *E. coli* ST10 and ST131 respectively.

### Oxford Nanopore Technologies (ONT) long-read sequencing

Serial isolates that developed non-CP-CRE, isolates used in the serial passaging experiments, and non-CP-CRE isolates from a previous study (18) underwent Oxford Nanopore Technologies (ONT) long-read sequencing. Library preps were completed using the SQK-RBK004 rapid barcoding kit with ~400 ng of input DNA and run on ONT MinION R9.4.1 flow cells using the ONT GridION X5 (Oxford, UK) per manufacturer’s instructions. ONT fast5 data was generated using ONT MinKNOW software (v3.0.13) with subsequent base-calling using Guppy v3.2.2 software (Oxford, UK). qcat-v1.1.0 was used for read demultiplexing, read length filtering (>1000 bp), and barcode removal (nanoporetech qcat GitHub: https://github.com/nanoporetech/qcat). A custom python script was used for the generation of polished, consensus assemblies (Shropshire, W flye_hybrid_assembly_pipeline GitHub: https://github.com/wshropshire/flye_hybrid_assembly_pipeline). Briefly, the Flye-v2.5 (24) assembler was used for *de novo* assembly, and contigs were circularized using berokka-v0.2 (Seemann, T berokka GitHub: https://github.com/tseemann/berokka). The circlator v1.5.5 (25) ‘clean’ command was then used to remove duplicate contigs sharing at least 90% identity. The remaining contigs were polished with Racon-v.1.4.5 (26) using the Oxford Nanopore long reads and then re-oriented with Circlator (25) “fixstart” to standardize the chromosome to the *dnaA* gene. A second long-read polish was then performed with Racon-v1.4.5 and these polished, corrected contigs were used as input for Medaka v0.8.1 long-read polishing (nanoporetech medaka GitHub: https://github.com/nanoporetech/medaka) followed by multiple rounds of Illumina short-read polishing using Racon-v1.4.5 (26). ONT assembly metrics are provided in **Table S1**.

### Phylogenetics, genetic variant calling, and clonality analysis

Phylogenetic analysis, *in silico* MLST, and subsequent variant-calling of serial isolates compared against the index strain ONT consensus assembly were used to determine clonality. Initially, the pan-genome pipeline tool Roary-v3.12.0 (27) was used to perform a core genome alignment of the short-read SPAdes assemblies using the probabilistic alignment program PRANK (27, 28). A core genome pairwise SNP distance matrix was generated with these data using a custom Python script (Narechania, A GitHub:https://github.com/narechan/amnh/blob/master/bin/snp_matrixBuilder.pl), which was subsequently used to build a maximum likelihood (ML) phylogenetic tree using RAxML-v8.2.12 (29). *In silico* multi-locus sequencing typing (MLST) was performed on the short-read SPAdes assemblies using mlst-v2.15.1 (Seemann, T mlst Github: https://github.com/tseemann/mlst; (30). Recurrent isolates that became non-CP-CRE were checked for clonality using the variant calling pipeline tool Snippy-v4.3.6 (Seemann, T snippy GitHub: https://github.com/tseemann/snippy) with INDELs removed.

### Genome annotation, AMR gene, and MGE identification

Gene calling and functional annotation was performed using Prokka-v1.14.0 (31). Annotated consensus genomes, as well as individual contigs, were parsed with ABRicate (Seemann, T ABRicate Github: https://github.com/tseemann/abricate) using the Comprehensive Antibiotic Resistance Database (CARD) (32) and PlasmidFinder (33) to search for AMR determinants and plasmid signatures respectively. Additionally, annotated Prokka gbk files were used to characterize ORFs of interests as well as confirm the results found with ABRicate. CARD, PlasmidFinder, ISFinder (34) and BLAST webtools were utilized during manual inspection of the assemblies to ensure correct context of annotated features, identify inverted repeat regions and target site duplications of insertion sequences, and confirm likely AMR gene mutations if present.

In cases where assemblies were not completely resolved due to putative large repeat regions that could not be fully captured on ONT long reads, we used a newly developed tool SVAnts-v0.1(35) (Hanson, Blake SVAnts Github: https://github.com/EpiBlake/SVAnts) to investigate subsets of ONT long reads. This tool enabled us to find individual ONT long reads containing MGEs of interest and align the DNA bordering these MGEs to our index assemblies to identify the regions in which the MGE was inserted within each bacterial genome.

### B-lactamase encoding gene and gene transcript level analysis

Quantitative PCR (qPCR) and quantitative real-time PCR (qRT-PCR) was used to assess both DNA copy number and RNA transcript levels respectively. Strains were grown in triplicate on two separate days (six biologic replicates) to mid-exponential phase (OD_600_ ~ 0.5) in Luria-Bertani (LB) broth (ThermoFisher) at 37° C shaking at 220 rpm. DNA isolation was performed using the DNEasy kit (Qiagen) and qPCR was performed using TaqMan reagents on the StepOne Plus Real Time PCR platform (Applied Biosystems). The DNA levels of *bla*_OXA-1_ and *bla*_CTX-M_ were determined relative to the *rpsL* control gene using the ΔCt method (6).

For RNA transcript level analysis, cells were mixed 1:2 with RNAProtect (Qiagen) and harvested via centrifugation. RNA was isolated from cell pellets using the RNEasy kit (Qiagen) and converted to cDNA using the High Capacity cDNA Reverse Transcription Kit (Applied Biosystems). Relative transcript levels of the β-lactamase encoding genes (*bla*_OXA-1_ and *bla*_CTX-M_) were assayed using TaqMan reagents on the StepOne Plus Real Time PCR machine (Applied Biosystems). The transcript level of *bla*_OXA-1_ and *bla*_CTX-M_ were determined relative to the endogenous control gene *rpsL* (36) using the ΔCt method. qPCR and qRT-PCR primers and probes are provided in **Table S2**.

### Serial passaging experiments on clinical *E. coli* ST131 index bacteremia isolate

p4A passaging experiments with antibiotic selection were performed in LB broth under shaking conditions at 37°C. A single colony from the index strain of patient 4 (p4A) was grown overnight, then diluted 1:100 into fresh LB containing ertapenem (ETP) (Sigma-Aldrich) at 0.5 MIC ETP. The process was repeated with increasing concentrations of ETP until growth was observed at an ETP concentration of 32 μg/mL. Passaging experiments were performed twice with growth occurring at an ETP MIC of ≥ 32 μg/mL within 3 passages (i.e. 72 hours) on each occasion. For the first passaging experiment, cells were collected on four consecutive days (strains p4A_1 to p4A_4) while continuing to passage at an ETP concentration of 32 μg/mL. This was done in order to determine whether progressive β-lactamase encoding gene amplification would be observed. On the second set of passaging experiments, cells were collected on the first day that the ETP MIC reached ≥ 32 μg/mL, serially diluted onto agar plates, and two different isolates were studied to assess for heterogeneity (strains p4A_H1 and p4A_H2). Additional serial passaging experiments were conducted using p4C and p4D isolates in the absence of antibiotic selection to determine the stability of the amplified units with protocol adapted from previously published methods (37).

### β-lactamase cloning and expression analysis

The open reading frames of β-lactamases were amplified from genomic DNA of the *K. pneumoniae* strain MB101 using Q5 polymerase and the primers listed in **Table S2**. Cloned ORFs were inserted into the arabinose inducible vector pBAD33 by Gibson assembly (38) of purified products. pBAD33*bla*_CTX-M-15_ and pBAD33*bla*_OXA-1_ were transformed into DH5α *E. coli* and were maintained with 50 μg/mL chloramphenicol in cation adjusted Mueller Hinton (MHII) media. MIC assays were performed with ceftriaxone (Sandoz GmbH), or piperacillin-tazobactam (Fresenius Kabi USA) as follows. DH5α strains carrying pBAD33*bla*_CTX-M-15_, pBAD33*bla*_OXA-1_, or control vector were grown for 18 hours at 220 rpm at 37°C in MHII with 50 μg/mL chloramphenicol. These cultures were diluted to approximately 5×10^5^ CFU/mL in MHII with 0.2% L-Arabinose or vehicle control, without chloramphenicol. Each strain was exposed to serial dilutions of the above drugs in microtiter plates sealed with gas-permeable membranes (Midsci). Microtiter plates were incubated for 18 hours at 220 rpm at 37°C, followed by OD_600_ measurement in a Biotek Synergy HT plate reader. The lowest tested antibiotic concentration yielding OD_600_ measurement of 0.06 or less in at least two of three replicate wells was considered to be the MIC.

### Statistical analyses

All statistical analyses were performed using Stata v13.1 (StataCorp LP, College Station, TX). Bivariate comparisons between patients with recurrent bacteremia and patients with a single bacteremia episode were made with the Wilcoxon Rank-sum test and Fisher’s exact test as appropriate based on covariate distributions. Comparisons of DNA and RNA levels among strains was performed using the Kruskall-Wallis test when more than two strains were analyzed or the Wilcoxon Rank-sum test when two strains were analyzed. MIC comparisons were performed using ANOVA with Dunnett’s test of multiple comparisons. Statistical significance was assigned as a two-sided *P* value < 0.05.

### Data availability

The index isolate assemblies from recurrent bacteremia patients that developed non-CP-CRE as well as the long-read and short-read data for all isolates have been deposited in the National Center for Biotechnology (NCBI) BioProject database PRJNA603908. ONT-sequencing data of non-CP-CRE isolates from a previous study (18) were deposited in NCBI BioProject database PRJNA388450. All other data analyzed during this study are available in the supplemental materials and/or available upon request from the corresponding author.

## RESULTS

### 116 patients with ESBL-E bacteremia were identified from the University of Texas MD Anderson Cancer Center (MDACC) from January 2015 to July 2016

Clinical and demographic features are presented on **Table S3**. *E. coli* was the most common organism isolated (100/116; 86.2%), followed by *K. pneumoniae* (14/116; 12.1%), and *K. oxytoca* (2/116; 1.7%). Carbapenems were used as primary treatment in 92% of index cases. Recurrent bacteremia was identified in 16/116 (13.8%) patients and primarily occurred either in patients with leukemia or recipients of hematopoietic stem cell transplants (14/16 cases, **Table S3**). The majority (14/16) of recurrent bacteremia cases had *E. coli* isolated, with the remaining two cases being due to *K. pneumoniae* infections. Carbapenem-resistant isolates were present in 4/16 (25%) recurrent bacteremia patients (**Fig. 1**). All four recurrent isolates that developed carbapenem resistance were *E. coli* isolates from leukemia patients. The full set of serial isolates was available for 11/16 (68.8%) patients, including all strains from the four patients that had at least one recurrent isolate that developed a carbapenem resistant phenotype.

**FIG 1.**
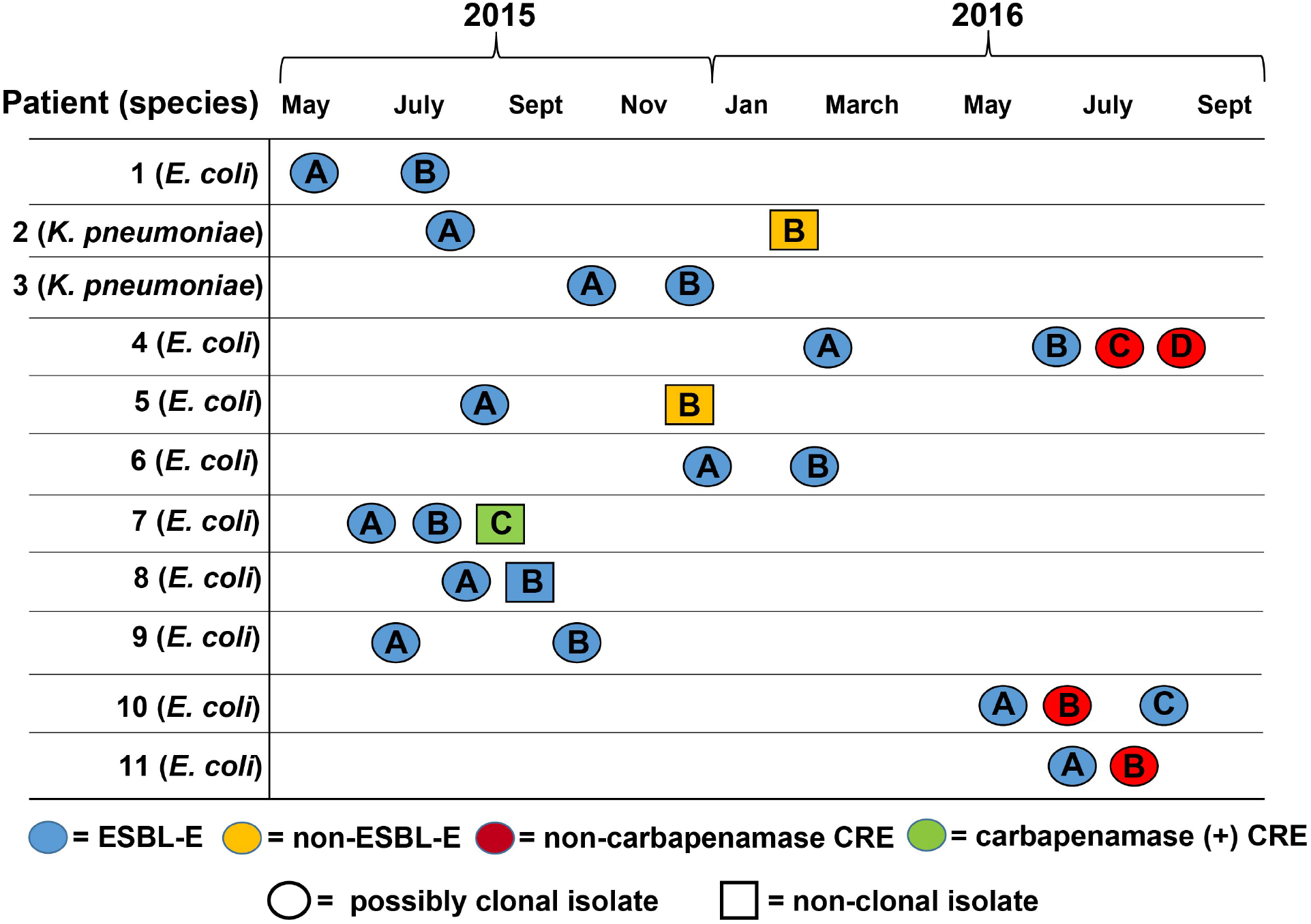
Overview of strains using Illumina short-read data. Timeline showing date of serial isolation from blood cultures. Patient numbers are in the first column. The shape and color of the isolates as labelled in the legend indicate clonality and antimicrobial resistance respectively. Strains were considered possibly clonal if they were the same sequence type and clustered on the phylogenetic tree. Patient subgroups (e.g. Patient 1; isolate A and B) refers to the order of isolation. Abbreviations are as follows: ESBL-E = extended spectrum β-lactamase producing *Enterobacterales*, CRE = carbapenem resistant *Enterobacterales*.

### Detection of non-CP-CRE emergence from three sets of clonal, ESBL-E recurrent bacteremia isolates

The 11 sets of patient serial isolates included nine *E. coli* serial isolates and two *K. pneumoniae* serial isolates. We performed Illumina short-read sequencing on the 26 strains identified from the 11 sets of serial isolates. Strain details are provided in **Table S4**. Serial strain relatedness was assessed using phylogenetic analysis with strains considered possibly clonal if they were the same sequence type, had the same Bayesian population structure, and clustered together on the phylogenetic tree (**Fig. S1**). The temporal collection, antimicrobial resistance, and strain relatedness are depicted in **Fig. 1**. Only a single strain, the 3^rd^ isolate from patient 7, which we will abbreviate as p7C, had a carbapenemase based on a positive Neo-Rapid CARB Kit Test result (**Fig. 1**). Whole genome, short-read sequencing analysis confirmed the presence of the class D carbapenemase, *bla*_OXA-1_81 in p7C. Furthermore, p7C was a different sequence type from the patient 7 index strain (p7A) indicating new strain acquisition.

We focused our subsequent analyses on the isolates from patient 4 (p4), patient 10 (p10), and patient 11 (p11), as each had at least one carbapenem resistant recurring non-CP-CRE isolate. Our HiSeq WGS data indicated that all three patients had recurrent *E. coli* isolates that respectively clustered together within a core genome phylogenetic tree, had the same Bayesian hierarchical population structure, and belonged to the same sequence type (**Fig. S1**). We confirmed clonality by measuring pairwise SNP distances between each recurrent isolate and their respective index strain using their highly resolved, ONT consensus assemblies. Our analysis indicated that there were less than 20 SNPs for all respective recurrent strains relative to their index strain suggesting that these three patients had clonal, re-infecting strains that had developed carbapenem resistance through a non-carbapenemase mechanism (**Table 1**).

**Table 1.**
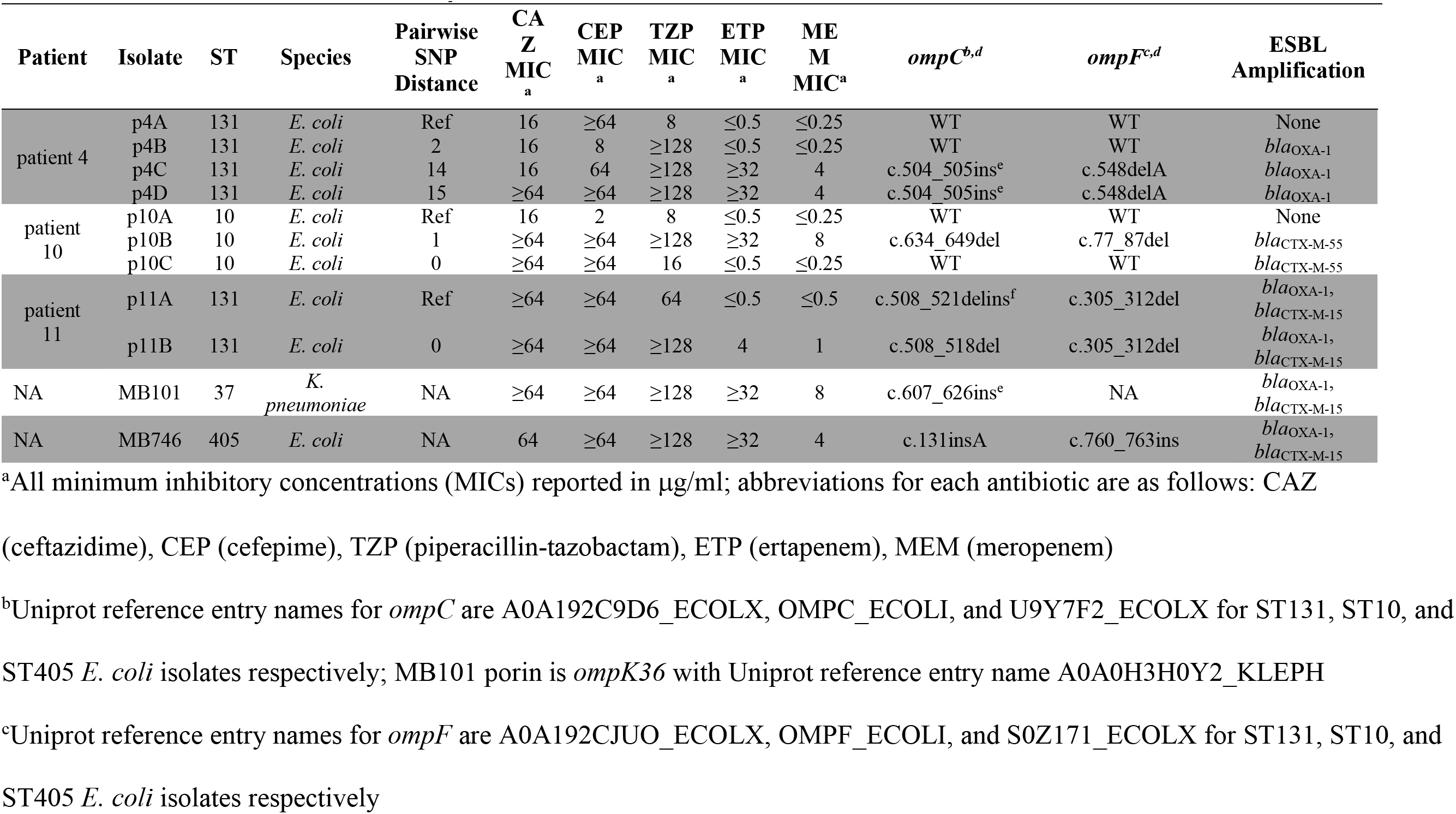
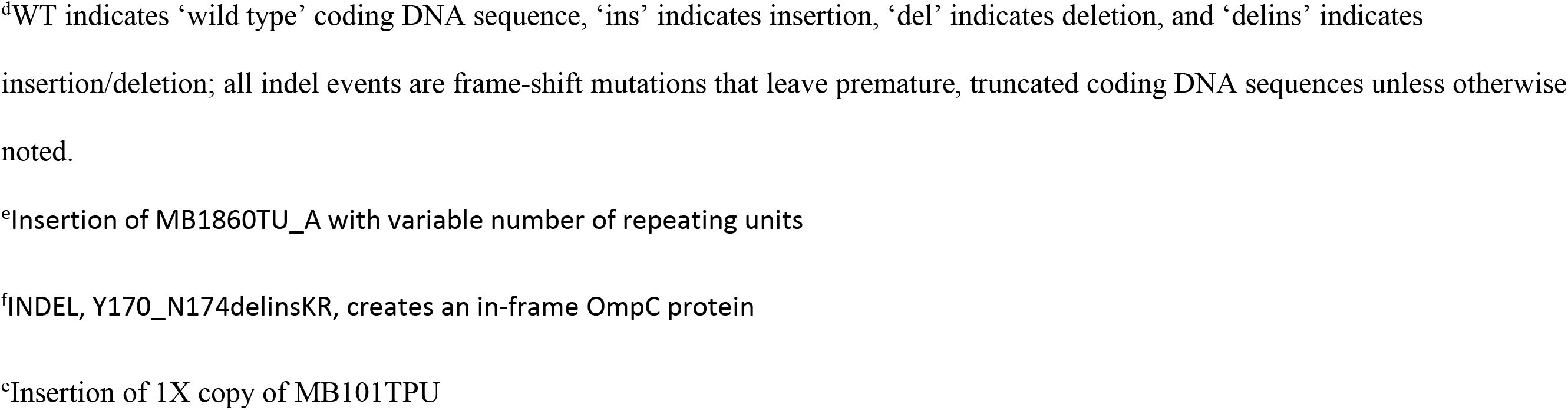
Clonality, Antibiotic Minimum Inhibitory Concentration (MIC), Outer Membrane Protein, and β-lactamase Characterization of *Enterobacterales* Isolates.

| Patient | Isolate | ST | Species | Pairwise<br>SNP<br>Distance | CA<br>Z<br>MIC<sub>a</sub> | CEP<br>MIC<sub>a</sub> | TZP<br>MIC<sub>a</sub> | ETP<br>MIC<sub>a</sub> | ME<br>M<br>MIC<sup>a</sup> | *ompC*<sup>b,d</sup> | *ompF*<sup>c,d</sup> | ESBL<br>Amplification |
| --- | --- | --- | --- | --- | --- | --- | --- | --- | --- | --- | --- | --- |
| patient 4 | p4A | 131 | *E. coli* | Ref | 16 | ≥64 | 8 | ≤0.5 | ≤0.25 | WT | WT | None |
|  | p4B | 131 | *E. coli* | 2 | 16 | 8 | ≥128 | ≤0.5 | ≤0.25 | WT | WT | *bla*<sub>OXA-1</sub> |
|  | p4C | 131 | *E. coli* | 14 | 16 | 64 | ≥128 | ≥32 | 4 | c.504_505ins<sup>e</sup> | c.548delA | *bla*<sub>OXA-1</sub> |
|  | p4D | 131 | *E. coli* | 15 | ≥64 | ≥64 | ≥128 | ≥32 | 4 | c.504_505ins<sup>e</sup> | c.548delA | *bla*<sub>OXA-1</sub> |
| patient 10 | p10A | 10 | *E. coli* | Ref | 16 | 2 | 8 | ≤0.5 | ≤0.25 | WT | WT | None |
|  | p10B | 10 | *E. coli* | 1 | ≥64 | ≥64 | ≥128 | ≥32 | 8 | c.634_649del | c.77_87del | *bla*<sub>CTX-M-55</sub> |
|  | p10C | 10 | *E. coli* | 0 | ≥64 | ≥64 | 16 | ≤0.5 | ≤0.25 | WT | WT | *bla*<sub>CTX-M-55</sub> |
| patient 11 | p11A | 131 | *E. coli* | Ref | ≥64 | ≥64 | 64 | ≤0.5 | ≤0.5 | c.508_521delins<sup>f</sup> | c.305_312del | *bla*<sub>OXA-1</sub>,<br>*bla*<sub>CTX-M-15</sub> |
|  | p11B | 131 | *E. coli* | 0 | ≥64 | ≥64 | ≥128 | 4 | 1 | c.508_518del | c.305_312del | *bla*<sub>OXA-1</sub>,<br>*bla*<sub>CTX-M-15</sub> |
| NA | MB101 | 37 | *K. pneumoniae* | NA | ≥64 | ≥64 | ≥128 | ≥32 | 8 | c.607_626ins<sup>e</sup> | NA | *bla*<sub>OXA-1</sub>,<br>*bla*<sub>CTX-M-15</sub> |
| NA | MB746 | 405 | *E. coli* | NA | 64 | ≥64 | ≥128 | ≥32 | 4 | c.131insA | c.760_763ins | *bla*<sub>OXA-1</sub>,<br>*bla*<sub>CTX-M-15</sub> |

<sup>a</sup>All minimum inhibitory concentrations (MICs) reported in µg/ml; abbreviations for each antibiotic are as follows: CAZ
(ceftazidime), CEP (cefepime), TZP (piperacillin-tazobactam), ETP (ertapenem), MEM (meropenem)
<sup>b</sup>Uniprot reference entry names for *ompC* are A0A192C9D6\_ECOLX, OMPC\_ECOLI, and U9Y7F2\_ECOLX for ST131, ST10, and
ST405 *E. coli* isolates respectively; MB101 porin is *ompK36* with Uniprot reference entry name A0A0H3H0Y2\_KLEPH
<sup>c</sup>Uniprot reference entry names for *ompF* are A0A192CJUO\_ECOLX, OMPF\_ECOLI, and S0Z171\_ECOLX for ST131, ST10, and
ST405 *E. coli* isolates respectively
<sup>d</sup>WT indicates ‘wild type’ coding DNA sequence, ‘ins’ indicates insertion, ‘del’ indicates deletion, and ‘delins’ indicates insertion/deletion; all indel events are frame-shift mutations that leave premature, truncated coding DNA sequences unless otherwise noted.
<sup>e</sup>Insertion of MB1860TU\_A with variable number of repeating units
<sup>f</sup>INDEL, Y170\_N174delinsKR, creates an in-frame OmpC protein
<sup>e</sup>Insertion of 1X copy of MB101TPU

### Gene amplification and porin loss associated with emergence of non-CP-CRE

Each of the serial isolates that developed a non-CP-CRE phenotype (p4A-D, p10A-C, and p11A-B) had increased Illumina short-read and ONT long-read coverage depth for β-lactamase encoding genes indicating amplification (**Fig. 2, Table S5 and Table S6**). Specifically, we noted increased coverage depth of *bla*_OXA-1_ in all p4 recurrent isolates, *bla*_CTX-M-55_ in p10 recurrent isolates, and *bla*_OXA-1_and *bla*_CTX-M-15_ in both the index and recurrent isolate of p11 (**Table 1**). We used qPCR to confirm the increased short-read and long-read coverage depth of our WGS analyses and found that the increased coverage depth corresponded to increased transcript levels of the amplified β-lactamase encoding genes (**Fig. 2**).

**FIG 2.**
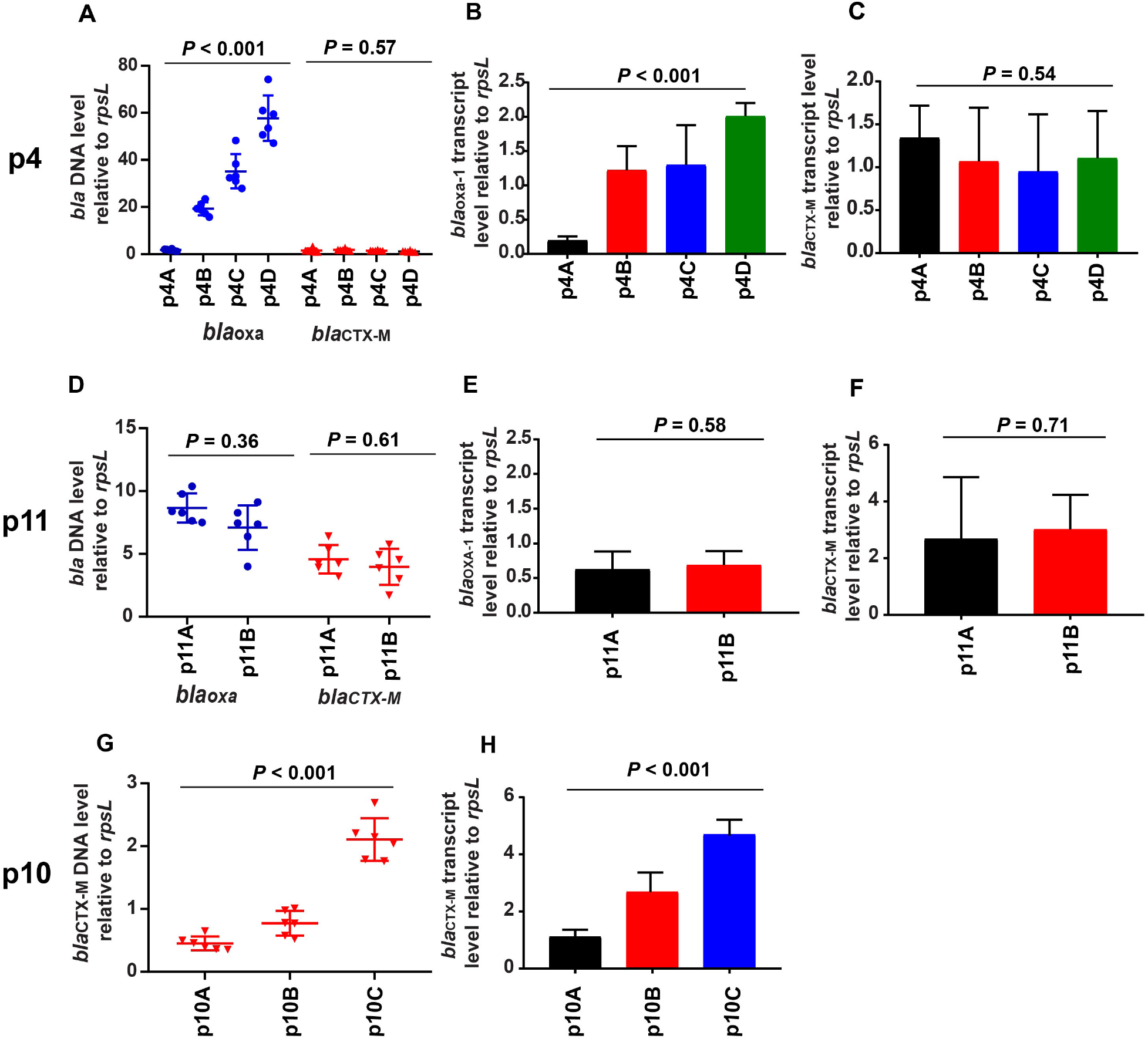
PCR analysis of β-lactamase encoding gene levels and transcript levels. Rows from top to bottom are data from patient 4 (p4), patient 11 (p11), and patient 10 (p10). (A, D, G) Taq-Man qPCR of genomic DNA results collected in triplicate on two separate days (n = 6) for either *bla*_OXA-1_ (blue) or *bla*_CTX-M_ (red) relative to the endogenous control gene *rpsL*. Data shown are individual data points with mean ± SD superimposed. (B, E, H) *bla*_OXA-1_ transcript level relative to endogenous control gene *rpsL*. RNA was collected from mid-exponential phase in triplicate on two separate days (n = 6). Data shown are mean ± SD. (C, F) similar analysis to (B, E, H) except that data are for *bla*_CTX-M_. Note that the p10 strains do not contain *bla*_OXA-1_. *P* values refer to measurements in the serial isolates relative to initial isolate using the Kruskal-Wallis (p4 and p10) or Wilcoxon rank-sum tests (p11).

We further characterized outer membrane porin genes with our sequencing data as disruptions of these genes are often correlated with carbapenem resistance. The only consistently identified variation in the non-CP-CRE strains relative to their ESBL-E index strains were mutations that disrupted the ORFs of the porin proteins OmpC and OmpF (**Table 1**). The initial sequencing results indicated that an insertion sequence (IS) as well as nucleotide deletions that resulted in frame-shifts mediated *ompC* gene disruption in isolates that developed non-CP-CRE. Conversely, we consistently found frame-shift inducing deletions in *ompF* genes for both the carbapenem susceptible and resistant isolates (**Table 1**). For p10 isolates, interruption of OmpC and OmpF in the second, non-CP-CRE isolate (p10B) was followed by reversion to the WT OmpC and OmpF genes in the third, carbapenem-susceptible isolate (p10C).

### Class 1 transposon *TnMB1860* found in both ST131 p4 and p11 serial isolates

The consensus ONT assembly index strains for patient 4 (p4A) and patient 11 (p11A) both contain an 8,147 bp, IS*26*-mediated translocatable unit (TU), designated as MB1860TU_A, which carries *bla*_OXA-1_, and inserts upstream adjacent to another 3,985 bp TU, designated MB1860TU_B, that carries *bla*_CTX-M-15_ (**Fig. 3A**). MB1860TU_C is the combination of both respective TUs that is 12,132 bp in length. The entire putative class 1 transposon structure designated Tn*MB1860* is 12,952 bp (**Fig. 3A**).

**FIG 3.**
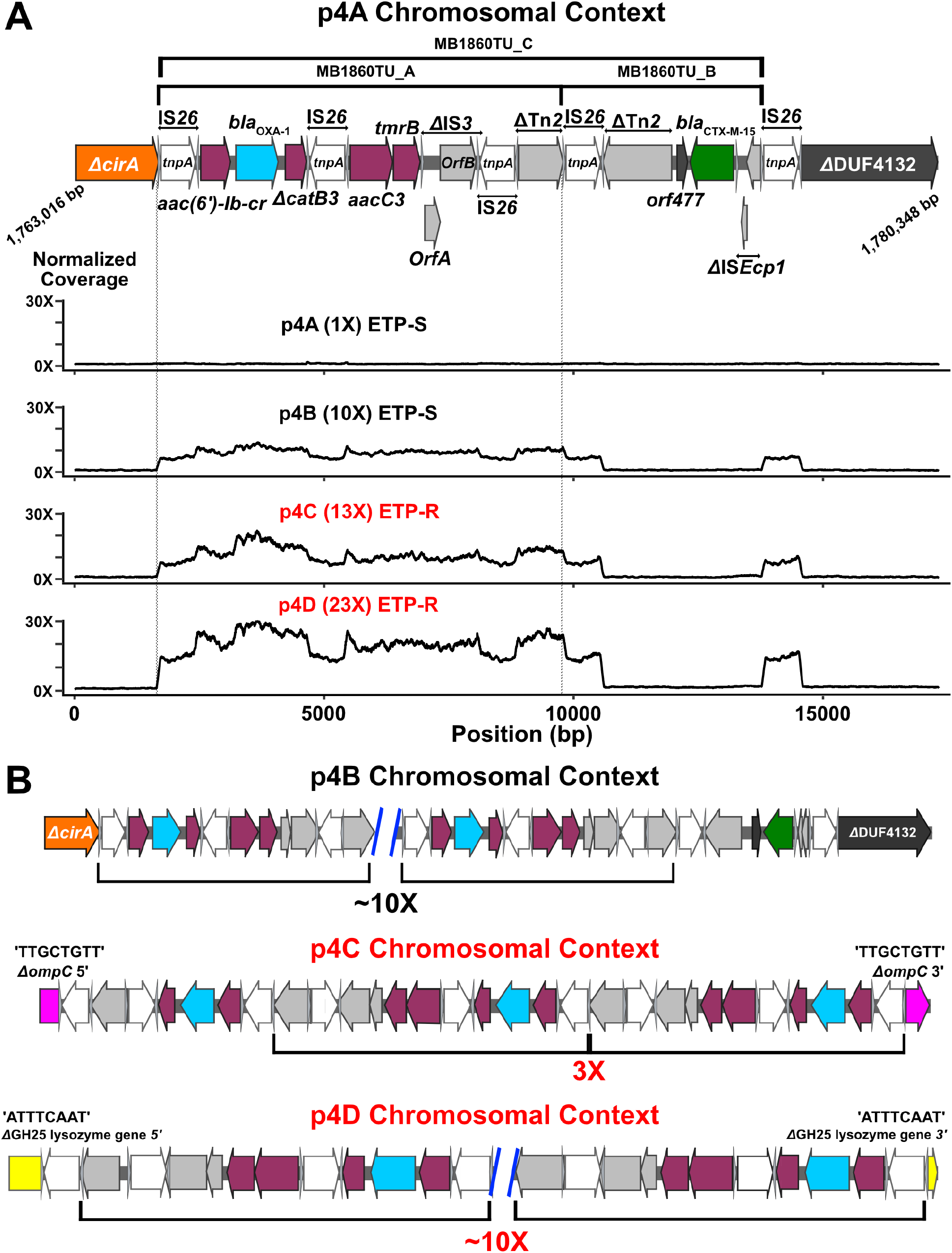
Characterization of IS*26*-flanked composite transposon with amplification and transposition of modular translocatable units in patient 4 serial isolates (i.e., p4A – p4D). Terminal left and right inverted repeats (IR_L_ and IR_R_ respectively) of insertion sequences (ISs) are specified by grey triangles that bracket respective complete and incomplete *tnpA* genes. ORFs are colored as follows: non-β-lactam AMR encoding genes (maroon), *bla*_OXA-1_ (blue), *bla*_CTX-M-15_ (green), IS*26 tnpA* (white), and other IS/Tn elements (gray). (A) Schematic indicates chromosomal context (1,763,016 – 1,780,348 bp; GenBank Accession #: CP049085)) of Tn*MB1860* locus flanked by directly oriented IS*26* transposases found in p4A isolate. Immediately below schematic are normalized, short-read coverage depth line graphs for the four p4 serial isolates with MB1860TU_A bracketed by dotted lines. ‘ETP-S’ = ertapenem susceptible; ‘ETP-R’ = ertapenem resistant. Font color for each serial isolate labelled in normalized coverage graph representing carbapenem susceptibility (black) or carbapenem resistance (red). (B) Characterization of amplification, transposition, and ORF disruption events in each of the respective patient 4 recurrent episode isolates. Black brackets beneath ORFs indicate MB168TU_A. The p4B chromosomal context shows a ~10X MB1860TU_A amplification event in the original p4A locus. The p4C chromosomal context additionally contains transposition and disruption of the *ompC* porin gene (pink) ~63 kbp upstream of original p4A *TnMB1860* locus with subsequent amplification. p4D isolate has previous amplification and transposition events found in p4B and p4C as well as another MB1860TU_A transposition and disruption of a putative glycoside hydrolase gene (i.e. GH25; yellow) ~71 kbp downstream of Tn*MB1860* locus with an amplification. The target site duplications (TSDs) created by the transposition of the TU are indicated above each respective junction site that flank the insertion and disruption of the OmpC and GH25 genes respectively.

When moving from the 5’ to 3’ end of Tn*MB1860* using the p4A chromosome (**Fig. 3A;** GenBank Accession #: CP049085) as the reference, the first resistance island contains two flanking IS*26 tnpA* genes in opposite orientation with the aminoglycoside N6’-acetyltransferase variant gene (*aac(6’)-Ib-cr*), the oxacillinase gene (*bla*_OXA-1_), and a truncated chloramphenicol resistance determinant (*ΔcatB3*). Following the second IS*26* element in Tn*MB1860*, there is an aminoglycoside N3’-acetyltransferase III variant gene, *aacC3,* and the tunicamycin resistance gene, *tmrB.* Downstream of the *tmrB* gene is an IS*3* element, which has a modified left inverted repeat (IR_L_) and a frame-shifted transposase that has been truncated by a third IS*26* element at the 3’ end of the IS*3* transposase. Immediately downstream of this IS*26* element is a truncated Tn2-like transposase which marks the 3’ boundary of MB1860TU_A. MB1860TU_A is inserted adjacent to the smaller translocatable unit, MB1860TU_B, which includes another fragment of a Tn2-like transposase, as well as *orf477*, *bla*_CTX-M-15_, and an IS*Ecp1* truncated by an intact IS*26*, which marks the 3’ boundary of both MB1860TU_B, as well as serves as the 3’ flanking IS*26* element for the full length Tn*MB1860*. Similar IS*26*/Tn2 family transposable elements that have putatively formed via homologous recombination events have previously been reported in association with IS*26*-mediated AMR transfer in *Enterobacterales* (6, 39, 40).

Tn*MB1860* is located on the p4 chromosome (1,764,660 – 1,777,611 bp; GenBank Accession #: CP049085) and the p11 chromosome (1,812,524 – 1,825,475 bp; GenBank Accession #: CP049077). One of the signatures of transposition are variable sized direct repeats called target site duplications (TSDs) that flank insertion sequences and are created during the transposition process (41–43). The Tn*MB1860* composite transposon on the p11A chromosome has 7-bp TSDs that indicate a transposition within a 3,762 bp ORF that putatively is involved in molybdopterin cofactor biosynthesis (**Fig. S2**). Interestingly, p4A differs in chromosomal context relative to p11A due to an intramolecular transposition event that occurred in reverse orientation (**Fig. S3)**. This is evidenced by the fact that p4A has an approximately 61 kbp region that is inverted with a downstream IS*26* in inverse orientation that has an 8-bp TSD within the colicin I receptor gene, *cirA*, in reverse complement orientation (**Fig. S3**). An alignment of p4A and p11A with two other *E. coli* ST131 chromosomes, TO217 (GenBank Accession #: LS992192.1) and Ecol_AZ146 (GenBank Accession #: CP018991.1) indicate similar chromosomal carriage of Tn*MB1860* with the noted inversion event that has occurred in the p4A isolate (**Fig. S3**).

### Delineation of MB1860TU_A amplification and transposition in ST131 p4 and p11 serial isolates

Patient 11 isolates p11A and p11B had relatively the same increase in mapping coverage for both *bla*_OXA-1_ and *bla*_CTX-M-15_ (**Table S3**). The consensus assemblies of both p11 isolates revealed that this increase in relative short-read coverage was due to two copies of *bla*_OXA-1_ and *bla*_CTX-M-15_ being present on a chromosomally located Tn*MB1860* as well as on IS*26*-mediated genomic resistant modules present on a 180,963bp multireplicon, F-type plasmid, p11A_p2 (**Fig. S2, Fig. S3;** GenBank Accession #: CP049079). The CRE phenotype of p11B relative to p11A appears to be driven by additional inactivation of the *ompC* gene given that the *ompF* gene is truncated in both strains (**Table 1**).

In contrast to the p11 isolates, the amplification and transposition of the modular, translocatable units that compose Tn*MB1860* in p4 was completely in a chromosomal context (**Fig. 3**). There was a consistent increase in short-read coverage depth (**Fig. 3A**) of the entire MB1860TU_A structure up to approximately 23-fold in p4D relative to the seven housekeeping pubMLST genes for ST131. We first analyzed strain p4B, which had developed resistance to piperacillin-tazobactam (TZP) but remained carbapenem susceptible (**Table 1**). In p4B, MB1860TU_A generated a ~10X tandem array *in situ* most likely through a conservative, IS*26*-mediated transposition mechanism or homologous recombination (**Fig. 3B**) (44, 45). We were unable to assemble the full-length tandem array due to limitations in read length size. However, we were able to use the SVAnts tool to identify thirty individual reads with two or greater number of tandem arrays of MB1860TU_A with 7 individual reads having 4X copies of *bla*_OXA-1_. The occurrence of the 10X TU amplification at the original Tn*MB1860* locus was confirmed by aligning the p4B long-reads to the reference chromosome p4A using SVAnts in conjunction with a short-read pileup analysis. Additionally, all outer membrane protein genes were WT and remained intact with p4B. Given that p4B had developed resistance to TZP relative to p4A (**Table 1**), we sought to determine whether overexpression of *bla*_OXA-1_ could drive TZP resistance. Inducing *bla*_OXA-1_ expression through cloning under an arabinose responsive promoter increased TZP MIC 6.8-fold relative to uninduced cells (**Fig. S4**).

Next, we examined the non-CP-CRE serial strains p4C and p4D. Interestingly, in both p4C and p4D a transposition and insertion of MB1860TU_A into *ompC* was present approximately 63 kbp upstream of the original *Tn*MB1860 chromosomal locus (**Fig. 3B**). This *ompC* gene disruption was confirmed through the identification of multiple p4C long-reads > 30 kbp that covered the full transposition site as well as identifying this insertion on the p4C and p4D incomplete chromosomal assemblies. We found two individual long reads that were able to span the full length of the MB1860TU_A array (3X copies) which disrupts *ompC* for p4C and confirmed the exact MB1860TU insertion location within *ompC* (c.504_505ins; **Table 1**). We also were able to identify 8-bp TSDs (‘5-TTGCTGTT-3’) at each end of the *ompC* insertion sites, which indicates MB1860TU_A replicative transposition. The p4D assembly and individual long-reads indicated a second MB1860TU_A transposition and insertion ~67 kbp downstream of the Tn*MB1860* within a GH25 lysozyme gene (**Fig. 3B**). This transposition event could be confirmed by 8-bp TSDs (5’-TTGCTGTT-3’) flanking the full insertion site (**Fig 3B**). A progressive increase in both DNA and RNA levels for *bla*_OXA-1_ but not *bla*_CTX-M-15_ in the p4 serial isolates was confirmed using qPCR and qRT-PCR, respectively (**Fig. 2A-C**).

### Serial passage of patient 4 index strain with ETP elicits *bla*_OXA-1_ and *bla*_CTX-M-15_ amplification through unique translocatable units relative to *in vivo* recurrent strains

We passaged strain p4A in increasing concentration of ertapenem (ETP) to determine whether antimicrobial exposure was driving the genetic changes observed in our serial clinical isolates. Resistance to ETP developed within three passages corresponding to three days, upon which we collected strains for the next four days (p4A_1-p4A_4). Short-read and qPCR analyses demonstrated all four isolates had amplification of both *bla*_OXA-1_ and *bla*_CTX-M-15_ relative to the p4A index strain prior to ETP exposure (**Fig 4**). ONT sequencing on all four isolates indicated that similar to p4B, there was *in situ* amplification occurring at the original Tn*MB1680* chromosomal locus. However, the full-length TU, MB1860TU_C, which consists of MB1860TU_A and MB1860TU_B (**Fig 4A**) that harbor *bla*_OXA-1_ and *bla*_CTX-M-15_ respectively, was the amplifying structure for the passaged isolates in contrast to what we saw in the *in vivo* isolates where MB1860TU_A was the sole amplifying structure (**Table 2**).

**Table 2.** p4A Serial Passaging Characterization.

| Sample | ETP MIC (µg/ml) | *ompC* | *ompF* | *bla*<sub>OXA-1</sub> copy #<sup>a</sup> | *bla*<sub>CTX-M-15</sub> copy #<sup>a</sup> |
| --- | --- | --- | --- | --- | --- |
| p4A_1 | ≥ 32 | c.208T>C; p.Q70X | c.17_18insIS1A | 6 | 7 |
| p4A_2 | ≥ 32 | c.208T>C; p.Q70X | WT | 8 | 9 |
| p4A_3 | ≥ 32 | c.208T>C; p.Q70X | c.62_63insIS1A | 6 | 7 |
| p4A_4 | ≥ 32 | c.208T>C; p.Q70X | c.517_518insIS1A | 6 | 7 |
| p4A_H1 | ≥ 32 | IS1A variant mediated insertion 96 bp upstream of +1 *ompC* start site | WT | 1 | 9 |
| p4A_H2 | ≥ 32 | IS1A variant mediated insertion 96 bp upstream of +1 *ompC* start site | WT | 1 | 17 |

<sup>a</sup>See Materials and Methods for Calculation

**FIG 4.**
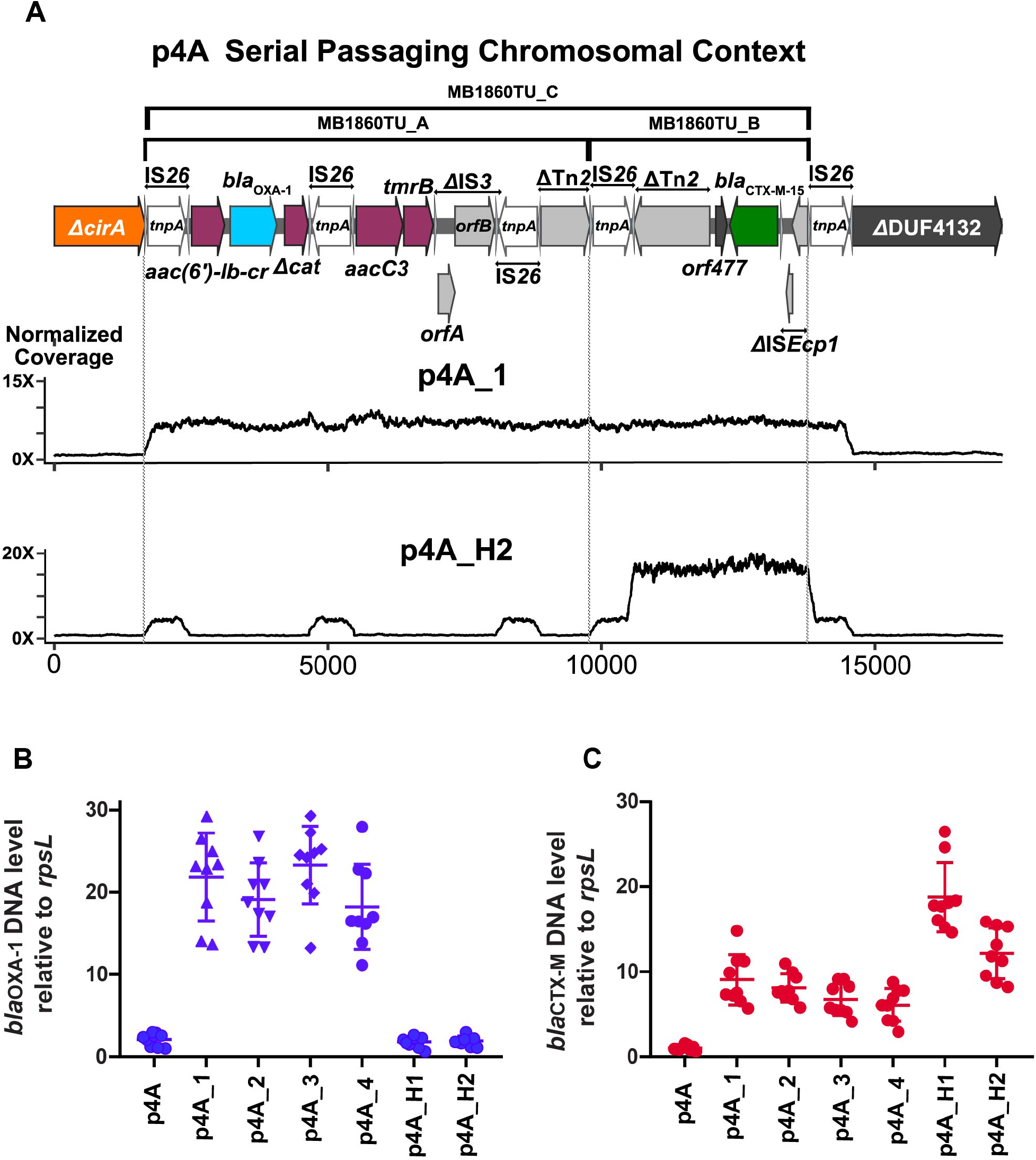
Identification and characterization of β-lactamase gene amplification following p4A serial passaging under ertapenem (ETP) exposure. Strain p4A was grown in ETP with isolates p4A_1-4 collected during the 1^st^ round of passaging and p4A_H1 and p4A_H2 collected during the 2^nd^ round. (A) Schematic of Tn*MB1860* locus from p4A as detailed in **FIG 2**. Immediately below the schematic are normalized, short-read coverage depth line graphs for p4A_1 and p4A_H2 aligned to p4A with location of MB1860TU_A and MB1860TU_B bracketed by dotted lines. Note amplification of MB1860 TU_C in strain p4A_1 whereas only MB1860TU_B amplified in strain p4A_H2. (B) and (C) are Taq-Man qPCR of genomic DNA collected in triplicate on two separate days (n = 6) for either *bla*_OXA-1_ (B) or *bla*_CTX-M_ (C) relative to the endogenous control gene *rpsL*. Data shown are individual data points with mean ± SD superimposed.

We repeated the experiment and again found that ETP resistance developed within three passages from the p4A isolate. ONT sequencing of the 2^nd^ round of passaged isolates (p4A_H1 and p4A_H2) revealed amplification of MB1860TU_B that harbors *bla*_CTX-M-15_ solely, which we verified using qPCR (**Fig. 4**). Similar to what we observed *in vivo*, the serially passaged ETP resistant isolates contained inactivating mutations in *ompC* although we did not observe any MB1860TU mediated interruptions (**Table 2**). INDELs inactivating the *ompF* gene were observed in a fraction of the serial isolates suggesting *ompF* inactivation may not be necessary for the development of non-CP-CRE (**Table 2**). We found that p4A responded *in vitro* to ETP similarly to what was observed in our serial clinical isolates by amplification of modular MB1860TU elements with concomitant porin disruption. Nevertheless, there was differential amplification of IS*26* translocatable units demonstrating the modularity of these MGEs.

In order to determine the chromosomal stability of the MB1860TU tandem arrays in the absence of antibiotic selective pressure, both ertapenem resistant (ETP-R) strains p4C and p4D were passaged for 10 days (~60 generations) without supplemented ertapenem. Both ETP-R recurrent strains consistently maintained carbapenem resistance through 60 generations of growth (**Fig. S5**). The p4C and p4D strains had relative copy number decreases of *bla*_OXA-1_ from 33 to 18X (45% decrease) and 53 to 42X (21% decrease) respectively (**Fig. S5**). These results indicate that both adapted strains can have persisting carbapenem resistance with associated tandem arrays of amplified *bla*_OXA-1_ maintained in the absence of antibiotic exposure.

### Unique IS*Ecp1*-mediated plasmid transposition of *bla*_CTX-M-55_ in patient 10 serial ST10 *E. coli* isolates

Compared to the p4 and p11 strains, the serial isolates from patient 10 (p10A – p10C) contain a different MGE that putatively drives the amplification of the ESBL encoding gene, *bla*_CTX-M-55_. Index strain p10A harbors *bla*_CTX-M-55_ located on an IS*Ecp1* transposition unit designated EC215TPU that resides on a multireplicon FIB-FIC-FII plasmid (**Fig. 5; GenBank Accession #: CP049082**). ‘Transposition unit’ (TPU) is used here to distinguish from translocatable unit (TU) as they have different transposition mechanisms that are mediated by IS*Ecp1* and IS*26* respectively. There are three additional small plasmids present in p10A, one of which is a ColE1-like 6.8 kbp plasmid designated as p10A_p2 (GenBank Accession #: CP049083), that is highly comparable to pCERC1 (GenBank Accession #: JN012467) (46).

**FIG 5.**
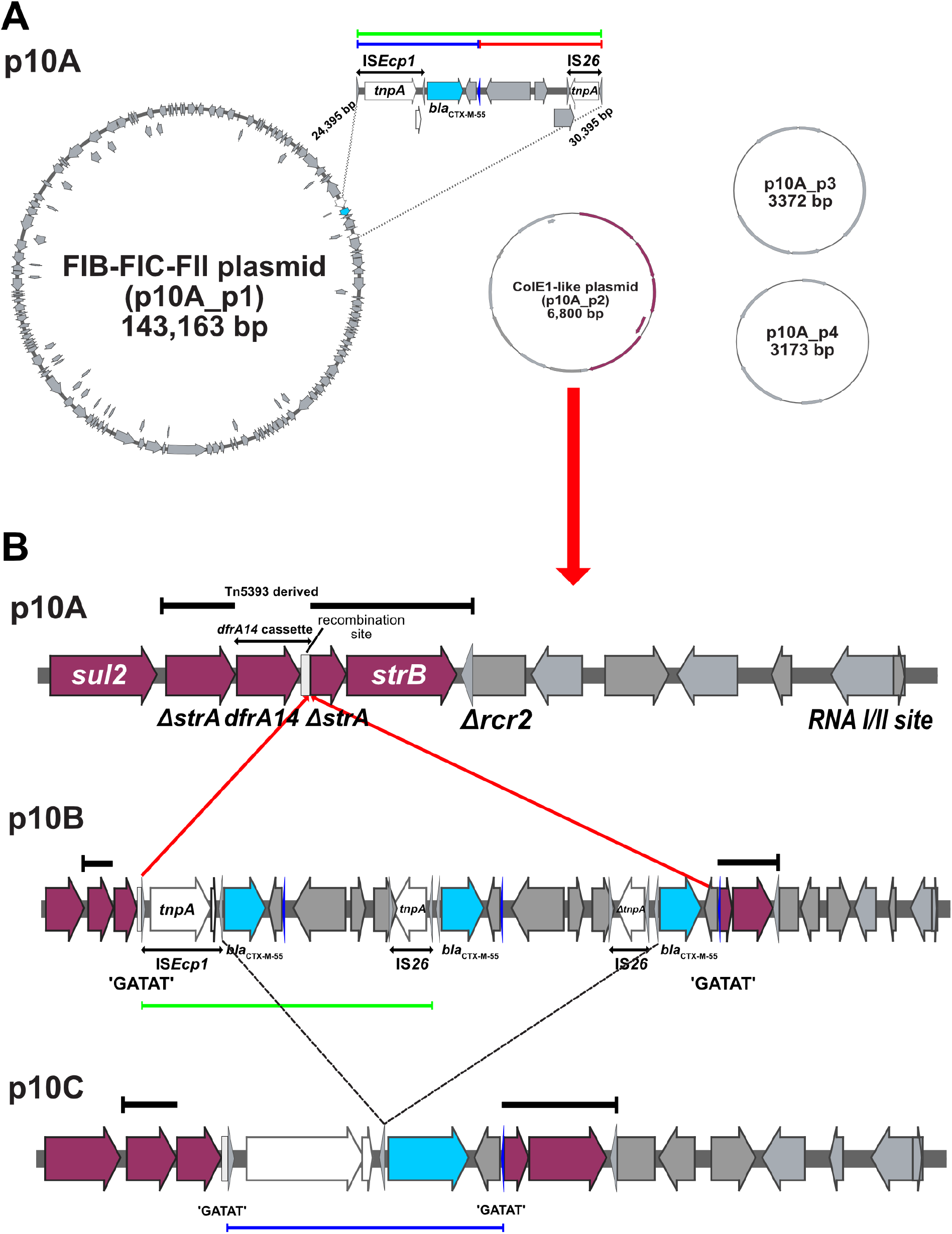
Genomic analysis of IS*Ecp1*-mediated mobilization of *bla*_CTX-M-55_ from a multireplicon F type plasmid (GenBank Accession #: CP049082) to a ColE1-like, high copy number plasmid (GenBank Accession #: CP049083) in p10 isolates. (A) Non-chromosomal genomic context for patient 10 index strain (p10A) harboring four circular plasmid structures. The *bla*_CTX-M-55_ gene is located on a 143,163 bp FIB-FIC-FII plasmid (p10A_p1) on an IS*Ecp1*-*bla*_CTXM-5-55_-IS*26* 5945 bp transposition unit (TPU) indicated by a green bracket. The blue and red regions are the union of this green bracketed region and represent regions highlighted in the following section. (B) Progression of TPU insertion into ColE1-like plasmid (p10A_p2). p10A illustrates the full genome context of p10A_p2 including the *sul2-ΔstrA-dfrA14-ΔstrA-strB* resistance island. The top black, bracketed region indicates genome that is derived from the Tn5393 transposon. The region indicated by double-ended arrows indicates where the *dfrA14* cassette inserted itself into Tn5393. The box downstream of *dfrA14* indicates the *attC* recombination site. p10B shows the amplified TPU (11,939 bp) insertion into the p10A_p2 recombination site (indicated by red arrows). The region is bracketed by 5-bp TSDs (5’-GATAT-3’) indicative of IS*Ecp1* mediated transposition. An alternative, right inverted repeat (IRRalt) is indicated by blue triangle. p10C contains an 8,998 bp deletion (black dotted lines) where two copies of *bla*_CTX-M-55_ are dropped. This creates a TPU flanked by the IRL and IRRalt of IS*Ecp1* (blue bracket).

EC215TPU translocates from the multireplicon, type F plasmid in p10A, designated p10A_p1, to p10A_p2 in the second serial isolate p10B (**Fig. 5B**). The IS*Ecp1*-mediated transposition can be identified and confirmed by the 5-bp TSDs that are immediately upstream of the IS*Ecp1* 14 bp IR_L_ and downstream of another alternative, 14 bp right inverted repeat (IRRalt) respectively (41, 42, 47). There are TSDs flanking EC215TPU at the p10A_p2 recombination site with both the IR_:L_ and IRRalt represented as a blue triangle (5’-CCTCACACCTTCGA-3’) on **Fig. 5B**. Notably, as the entire insertion region with three *bla*_CTX-M-55_ copies on p10B has 5 bp TSDs (5’-GATAT-3’) that flank IR_L_ and IR_Ralt_ respectively, one can postulate that an amplification event occurred on p10A_p1, which then subsequently inserted into p10A_p2 (**Fig. 5B)**. p10B is also the only carbapenem-resistant p10 serial isolate. This likely is due in part to deletions in *ompC* and *ompF* that create frame-shift truncations of each respective porin in contrast to the WT genotypes of each respective porin found in p10A and p10C. When analyzing the third isolate p10C, there only is one copy of *bla*_CTX-M-55_ identified in the assembly and short-reads on p10A_p2. However, the short-read and long-read coverage depth analysis suggests an increase in copy number (**Table S6**). The subsequent increase in initiation RNA genes, a marker for ColE1-like plasmids with *bla*_CTX-M-55_ suggests that p10C amplification occurs by an increase in the overall copy number of p10A_p2. These data are consistent with our qPCR analysis which demonstrate progressively higher DNA levels of *bla*_CTX-M-55_ for each serial isolate (**Fig. 2**).

### Detection of β-lactamase gene amplification and porin disruption by AMR elements in non-serial *Enterobacterales* strains

There were two non-CP-CRE strains, MB101 (*K. pneumoniae*) and MB746 (*E. coli*), identified in a previous study examining the role of short-read WGS in predicting β-lactam resistance (**Table 1)**(18). We performed ONT sequencing on the two isolates to determine whether mechanisms of gene amplification and porin disruption were similar to what was observed in the cohort of non-CP-CRE bacteremia serial isolates. Both isolates contain IS*26*-mediated TUs carrying *bla*_OXA-1_ as previously characterized in Tn*MB1860* (**Fig. 6**).

**FIG 6.**
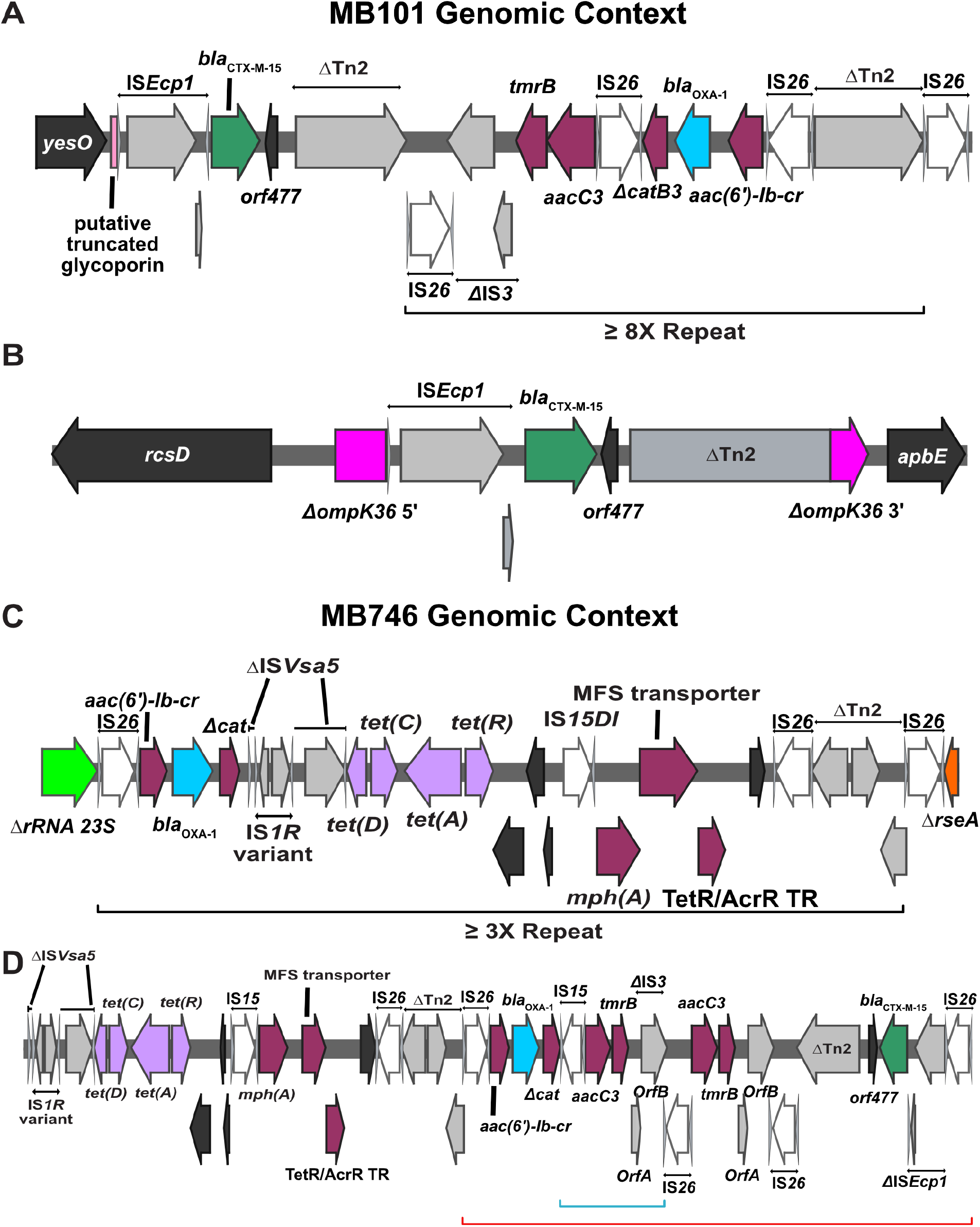
Genomic context of end-stage MB101 and MB746 isolates respectively. Terminal left andright inverted repeats (IR_L_ and IR_R_ respectively) of insertion sequences (ISs) are specified by grey triangles that bracket respective complete and incomplete *tnpA* genes. ORFs are colored as follows: AMR genes (maroon), *bla*_OXA-1_ (blue), *bla*_CTX-M-15_ (green), IS26 *tnpA* (white), and other IS/Tn elements (gray). Delta (Δ) next to annotated genetic region indicates a truncation or disruption (A-B) Chromosomal locations of MB101TU (Fig. 6A) and MB101TPU (Fig. 6B) indicating respective amplification and transposition of each element. ~8X MB101TU repeat indicated by black bracket. Truncated *ompK36* gene labelled in pink (Fig. 6B). (C-D) Genomic context of MB746TU and respective genomic resistance modules carrying AMR genes. Black brackets beneath Fig. 6C schematic indicate repeating MB101TU unit. Fig. 6D indicates FIB plasmid carriage of genomic resistance modules. Red bracket indicates IS*26* transposon structure that shares 100% coverage; 99% BLAST similarity with Tn*MB1860*. Blue bracket indicates small, 2X repeat structure.

For strain MB101 (**Fig. 6A-B**), a ST37 *K. pneumoniae* isolate, the ~9310 bp IS*26*-mediated TU, designated MB101TU harbors *aac-(6’)-Ib-cr, bla*_OXA-1_, Δ*catB3*, *aacC3,* and *tmrB* and is nearly identical (100% coverage; 99.9% ID) to MB1860TU_A found in serial *E. coli* isolates from patients 4 and 11 (**Fig. 6A**). An IS*Ecp1*-mediated TPU harboring *bla*_CTX-M-15_, designated MB101TPU, was also present on the MB101 chromosome as well as an FIB_K_ type plasmid, with at least five copies present on the chromosome based on the Flye assembly and individual long reads. The assembly and long reads indicate the transposition of MB101TPU into a putative glycoporin encoding gene which was followed by the transposition of MB101TU to an existing IS*26* element downstream of MB101TPU (**Fig. 6A**). We identified an amplification of MB101TU at this locus that created a tandem array of at least 8X copies based on the identification of multiple long-reads with multiple MB101TU copies. Interestingly, there was also a transposition event of MB101TPU into the *ompK36* encoding gene, a homolog of *ompC* in *E. coli* (**Fig. 6B**). Thus, MB101 has amplification of both *bla*_OXA-1_ and *bla*_CTX-M-15_ via distinct mechanisms along with TPU mediated porin disruption.

The ST405 *E. coli* isolate, MB746, has an IS*26*-mediated translocatable unit, designated MB746TU, which also includes a genomic resistance module that carries *aac(6’)-Ib-cr, bla*_OXA-1_, and a truncated chloramphenicol resistance determinant (**Fig. 6C-D**) similar to what is seen on MB1860TU_A. MB746TU carries an additional tetracycline resistance operon and macrolide resistance operon, not present on Tn*MB1860*. We found four individual long-reads that carried at least 3X copies of MB746TU present in the same MB746 chromosomal location (**Fig. 6C**). Additionally, we identified a FIB plasmid that harbored both *bla*_OXA-1_ and *bla*_CTX-M-15_ which has a similar configuration and orientation to Tn*MB1860* (Red bracket; **Fig. 6D**). The IS*26*-mediated TU carrying *bla*_CTX-M-15_ has 99.7% BLAST identity with MB1860_B with the only substantial difference being a size of 5558 bp vs the 3985 bp MB1860TU_B (**Fig. 6D**). Notably, MB746 also had *ompC* and *ompF* gene disruptions (**Table 1)**comparable to what was observed in serial isolates that developed non-CP-CRE phenotypes with concomitant β-lactamase amplifications.

These data show that there are multiple, non-CP-CRE clinical isolates within our region that have amplification and transposition of similar IS*26* and IS*Ecp1*-mediated elements that harbor β-lactamase genes encoding genes in conjunction with porin disruption.

## DISCUSSION

While recent systematic surveillance studies indicate that the prevalence of non-CP-CRE remains high (3, 4), the mechanisms contributing to non-CP-CRE emergence within clinical settings are largely unknown. This public health issue is particularly troublesome given the difficulties in properly diagnosing non-CP-CRE infections and prescribing effective antimicrobials due to the transient nature of their carbapenem resistance phenotypes as well as their lack of a carbapenemase. This analysis demonstrates that multiple TU- and TPU-mediated modular amplifications of β-lactamase encoding genes in conjunction with porin inactivation are driving non-CP-CRE emergence in our cohort of cancer patients with recurrent bacteremia . It is well established that “copy-in” replicative transposition is a common mechanism by which IS*26*-mediated TUs mobilize to regions lacking a pre-existing IS*26* element (43, 45, 48). Recent studies have shown that TUs can create tandem resistance gene arrays by targeting genomes with an existing copy of IS*26* through an intermolecular, conservative transposition mechanism and, less frequently, through RecA-dependent homologous recombination (43–45, 49). We were able to determine that the initial incorporation of the composite transposon Tn*MB1860* as well as MB1860TU_A transposition into *ompC* and the predicted glycoside hydrolase gene occurred through a replicative transposition mechanism based on characteristic TSDs at insertion sites. The amplification of MB1860TU_A, MB1860TU_B, and MB1860TU_C that created tandem arrays of resistance genes observed in our study likely occurred through a conservative transposition mechanism although RecA-dependent homologous recombination could also be contributing to the generation of these tandem arrays. The ability to detect genetic structures capable of amplifying β-lactamase encoding genes that synergistically insert into porin encoding genes will be essential for diagnostic and surveillance purposes in order to ensure we are properly identifying clinical isolates that have the ability to develop CRE.

Amplification of *bla*_OXA-1_ in both serial and non-serial isolates consistently included *aac-(6’)-Ib-cr* and Δ*catB3* in association with other modular, IS*26*-mediated translocatable units carrying resistance genes. Livermore et al. recently observed the co-occurrence of *aac-(6’)-Ib’* and *bla*_OXA-1_ in 147/149 *E. coli* strains, primarily ST131 isolates, suggesting that many *E. coli* isolates carrying *bla*_OXA-1_ may be capable of this amplification (40, 50). The finding that *bla*_OXA-1_ amplification was consistently associated with progressive development of β-lactam resistance was somewhat surprising given that this enzyme is typically considered a narrow spectrum β-lactamase (51). However, overexpression of *bla*_OXA-1_ using an arabinose inducing promoter was shown to generate resistance to ertapenem in a porin deficient strain (6). This study, along with our own results, indicates that *bla*_OXA-1_ likely contains sufficient carbapenem hydrolysis activity to generate resistance with the combination of augmented gene copy number and decreased carbapenem concentration due to porin inactivation. Additionally, we demonstrate that overexpression of *bla*_OXA-1_ without porin inactivation produced TZP resistance, a finding that could help resolve previously noted discrepancies between β-lactamase gene content and TZP susceptibility (18, 52). Similar amplifications of the narrow-spectrum, TEM β-lactamases have also been shown to be associated with TZP resistance, suggesting amplification of narrow-spectrum β-lactamases may be a substantial mechanism contributing to TZP resistance (9–11).

The other amplified β-lactamase in our cohort was *bla*_CTX-M_. CTX-M is the most commonly identified ESBL enzyme in *Enterobacterales* (53) and high level CTX-M production has previously been shown to confer carbapenem resistance in porin-deficient *E. coli* (6) as well as *K. pneumoniae* (54). Amplification of MB1860TU_B harboring *bla*_CTX-M-15_ with concomitant porin disruption was associated with development of ETP resistance in one of our two passaging experiments, demonstrating the versatility of various modules of Tn*MB1860* in responding to carbapenem exposure. Unlike *bla*_OXA-1_, we did not identify *bla*_CTX-M_ genes in association with other AMR encoding elements, but rather *bla*_CTX-M_ genes were consistently present with an IS*ECp1* element, which has been previously described for *E. coli* (55, 56). The array of mechanisms by which *bla*_CTX-M_ copy numbers could increase included (1) *in situ* IS*26*-mediated amplification of MB1860TU_B (strain P4A_H1) and MB1860TU_C (strain p4A1 – p4A4); (2) IS*Ecp1*-mediated transposition of EC215TPU from a low-copy, F-type plasmid to a multi-copy, ColE1-like plasmid (strain p10B); (3) and multiple chromosomal IS*Ecp1*-mediated transposition unit insertions including into the *ompK36* porin encoding gene (strain MB101). Thus, the various *bla*_CTX-M_ amplification mechanisms suggest that augmentation of *bla*_CTX-M_ copy number may be a major driver of non-CP-CRE development.

Along with permitting direct visualization of genomic resistance module amplifications containing various β-lactamase genes, ONT sequencing also allowed for identifying porin-mediated disruption by TU and TPU structures. Laboratory studies that generated CRE strains through serial passaging have consistently found that OmpC and OmpF porin production is reduced in carbapenem resistant isolates, but mechanisms have generally involved alterations in the porin regulatory protein OmpR or in OmpR binding sites (6, 7). Similarly, a recent study of a single patient with recurrent *E. coli* infection identified a single amino acid change in OmpR as leading to loss of OmpC and OmpF expression with development of ertapenem resistance (15). Conversely, we observed direct inactivation of OmpC and OmpF either through TU/TPU transposition into their respective ORFs or through INDELs that resulted in frame-shift mutations whereas no alterations in OmpR were identified in our cohort. To our knowledge, there have been only two other reports of an IS*Ecp1*-mediated TPU harboring a β-lactamase encoding gene causing the disruption of a porin, in both cases OmpK35 from *K. pneumoniae* (57, 58). While there have been a number of studies that have implicated IS-mediated porin disruptions, including IS*26* mediated disruptions (54, 59–61), to our knowledge, this is the first documented IS*26*-mediated translocatable unit disruption of an outer membrane porin followed by amplification of a β-lactamase gene. Given the length of the TUs, as well as the fact that they have the ability to amplify and create tandem arrays once inserted into the porin encoding genes, it is unlikely that targeted PCR based strategies or commonly used short-read approaches alone would have been able to identify the full extent of the TUs observed in our study. Thus, performing long read sequencing on larger cohorts of *Enterobacterales* with porin deficient backgrounds should reveal whether TU/TPUs harboring AMR genes are a frequent mediator of porin gene disruption, but have not been previously identified due to the long and repetitive nature of the involved DNA structures.

The clinical impact of β-lactamase gene amplification and porin loss driving carbapenem resistance has been postulated to be mitigated by the fitness costs imposed on the organism by such genetic changes (37, 62–64). However, such AMR mechanisms are increasingly being recognized as commonly occurring in clinical isolates, including in the serious infections described in our cohort (37, 62–64). This would suggest that organisms with the capability to amplify AMR genes are widespread and capable of causing significant human infections especially under antibiotic selective pressure (7, 9, 10, 65). Moreover, the most recent systematic data on CRE in the U.S. found no difference in outcomes between patients with CRE and non-CP-CRE suggesting that circulating non-CP-CRE organisms may not have significant fitness defects (4). Furthermore, our own results from serial passaging p4C and p4D without ETP exposure suggests these amplified structures harboring AMR genes associated with increased MICs to ertapenem and meropenem may be fairly stable in the absence of antibiotic selective pressure. The chromosomal context of Tn*MB1860* may provide insights into the stability of this structure as a previous study had noted stable transposon carriage when mobilized from plasmid to chromosome at very low levels of antibiotic exposure (66).

In addition to possibly imposing a fitness cost on the organism, AMR gene amplifications have also been associated with the presence of antimicrobial heteroresistance (65, 67). The interplay between AMR gene amplification, heteroresistance and fitness is likely reflected in the genotype changes of patient 10 isolates given the development and subsequent reversion of porin mutations observed in those strains (Table 1). Recent systematic surveys have indicated approximately 25% of strains reported as carbapenem resistant tested as susceptible at a central laboratory, likely reflecting the transient phenomenon of AMR heteroresistance (4, 64). A limitation of our study is that we only sequenced a single colony for each isolate, which almost certainly underestimates the genetic complexity of a bacterial population responding to antimicrobial therapy. Nevertheless, a challenge presents itself in capturing population heterogeneity with a single genome assembly as multiple, heterogeneous sequencing reads will break current assembler algorithms.

## CONCLUSIONS

We have used serial, clinical isolates subjected to complementary WGS approaches to identify that a combination of porin inactivation and IS-mediated amplification of TU and TPU elements harboring β-lactamase encoding genes underlies the emergence of non-CP-CRE from ESBL-E parental isolates within our study population. We predict that more widespread application of long-read sequencing technologies will facilitate appreciation of the mechanisms and impact of TU- and TPU-mediated transpositions, porin disruptions, and gene amplifications on a diverse array of AMR pathogens in the clinical setting.

## LIST OF ABBREVIATIONS

non-CP-CRE: non-carbapenemase-producing carbapenem resistant *Enterobacterales*
TU: translocatable unit
TPU: transposition unit

## DECLARATIONS

### Ethics approval and consent to participate

A waiver of informed consent to collect clinical data from electronic medical records and analyze the isolates was provided by the MDACC IRB (PA15-0799). A waiver of consent from UTHealth Science Center at Houston (IRB #: HSC-SPH-20-0032) was obtained to perform bacterial genomics sequencing and analysis.

### Consent for publication

Not applicable

### Author contributions

WCS developed the database, generated and analyzed the data, and was a major contributor to the writing of the manuscript. SLA designed the study, collected and analyzed the data, and significant contributor to writing the manuscript. RP helped designed the study, performed the experiments, and analyzed the data. JK performed experiments and analyzed the data. MB performed the experiments and contributed to writing the paper. XL helped perform phylogenomic analyses. AK helped perform phylogenomic analyses and contributed to writing the paper. JGP contributed to the design of the study, analyzed the isolates, and writing of the paper. PS collected and curated the isolates. CAA helped design the study, analyze the data, and writing of the paper. DEG analyzed the data and contributed to the writing of the paper. BMH wrote scripts to analyze the data, analyzed the data, and contributed to writing the paper. SAS designed the study, analyzed the data, and major contributor to the writing of the manuscript. All authors read and approved the final manuscript.

## Acknowledgements

We thank the personnel of the clinical microbiology laboratory at MD Anderson Cancer Center for assistance with collecting isolates. Would also like to thank Jennifer Walker, PhD for her insightful input in drafting the manuscript.

## REFERENCES

1. CDC. Antibiotic Resistance Threats in the United States, 2019. Atlanta, GA: U.S. Department of Health and Human Services, CDC; 2019.

2. Logan LK, Weinstein RA. The Epidemiology of Carbapenem-Resistant Enterobacteriaceae: The Impact and Evolution of a Global Menace. J Infect Dis. 2017;215(suppl_1):S28–S36.

3. Guh AY, Bulens SN, Mu Y, Jacob JT, Reno J, Scott J, et al. Epidemiology of Carbapenem-Resistant Enterobacteriaceae in 7 US Communities, 2012-2013. JAMA. 2015;314(14):1479–87.

4. Duin Dv, Arias C, Komarow L, Chen L, Hanson B, Weston G, et al. Molecular and Clinical Epidemiology of Carbapenem-Resistant Enterobacteriaceae in the United States: a Prospective Cohort Study. Lancet ID. 2020.

5. Tangden T, Adler M, Cars O, Sandegren L, Lowdin E. Frequent emergence of porin-deficient subpopulations with reduced carbapenem susceptibility in ESBL-producing Escherichia coli during exposure to ertapenem in an in vitro pharmacokinetic model. J Antimicrob Chemother. 2013;68(6):1319–26.

6. Adler M, Anjum M, Andersson DI, Sandegren L. Influence of acquired beta-lactamases on the evolution of spontaneous carbapenem resistance in Escherichia coli. J Antimicrob Chemother. 2013;68(1):51–9.

7. van Boxtel R, Wattel, A. A., Arenas, J., Goessens, W. H., & Tommassen, J. Acquisition of Carbapenem Resistance by Plasmid-Encoded-AmpC-Expressing Escherichia coli. Antimicrobial agents and chemotherapy. 2017;61(1):e01413–16.

8. Beceiro A, Maharjan S, Gaulton T, Doumith M, Soares NC, Dhanji H, et al. False extended-spectrum {beta}-lactamase phenotype in clinical isolates of Escherichia coli associated with increased expression of OXA-1 or TEM-1 penicillinases and loss of porins. J Antimicrob Chemother. 2011;66(9):2006–10.

9. Schechter LM, Creely DP, Garner CD, Shortridge D, Nguyen H, Chen L, et al. Extensive Gene Amplification as a Mechanism for Piperacillin-Tazobactam Resistance in Escherichia coli. MBio. 2018;9(2).

10. Rodriguez-Villodres A, Gil-Marques ML, Alvarez-Marin R, Bonnin RA, Pachon-Ibanez ME, Aguilar-Guisado M, et al. Extended-spectrum resistance to beta-lactams/beta-lactamase inhibitors (ESRI) evolved from low-level resistant Escherichia coli. J Antimicrob Chemother. 2019.

11. Hansen KH, Andreasen MR, Pedersen MS, Westh H, Jelsbak L, Schonning K. Resistance to piperacillin/tazobactam in Escherichia coli resulting from extensive IS26-associated gene amplification of blaTEM-1. J Antimicrob Chemother. 2019;74(11):3179–83.

12. Poirel L, Heritier C, Spicq C, Nordmann P. In vivo acquisition of high-level resistance to imipenem in Escherichia coli. J Clin Microbiol. 2004;42(8):3831–3.

13. Oteo J, Delgado-Iribarren A, Vega D, Bautista V, Rodriguez MC, Velasco M, et al. Emergence of imipenem resistance in clinical Escherichia coli during therapy. Int J Antimicrob Agents. 2008;32(6):534–7.

14. Chia JH, Siu LK, Su LH, Lin HS, Kuo AJ, Lee MH, et al. Emergence of carbapenem-resistant Escherichia coli in Taiwan: resistance due to combined CMY-2 production and porin deficiency. J Chemother. 2009;21(6):621–6.

15. Dupont H, Choinier P, Roche D, Adiba S, Sookdeb M, Branger C, et al. Structural Alteration of OmpR as a Source of Ertapenem Resistance in a CTX-M-15-Producing Escherichia coli O25b:H4 Sequence Type 131 Clinical Isolate. Antimicrob Agents Ch. 2017;61(5).

16. Kao CY, Chen JW, Liu TL, Yan JJ, Wu JJ. Comparative Genomics of Escherichia coli Sequence Type 219 Clones From the Same Patient: Evolution of the IncI1 blaCMY-Carrying Plasmid in Vivo. Front Microbiol. 2018;9:1518.

17. Zou H, Xiong SJ, Lin QX, Wu ML, Niu SQ, Huang SF. CP-CRE/non-CP-CRE Stratification And CRE Resistance Mechanism Determination Help In Better Managing CRE Bacteremia Using Ceftazidime-Avibactam And Aztreonam-Avibactam. Infect Drug Resist. 2019;12:3017–27.

18. Shelburne SA, Kim J, Munita JM, Sahasrabhojane P, Shields RK, Press EG, et al. Whole-Genome Sequencing Accurately Identifies Resistance to Extended-Spectrum beta-Lactams for Major Gram-Negative Bacterial Pathogens. Clin Infect Dis. 2017;65(5):738–45.

19. Cao MD, Nguyen SH, Ganesamoorthy D, Elliott AG, Cooper MA, Coin LJ. Scaffolding and completing genome assemblies in real-time with nanopore sequencing. Nat Commun. 2017;8:14515.

20. Harris PA, Taylor R, Thielke R, Payne J, Gonzalez N, Conde JG. Research electronic data capture (REDCap)--a metadata-driven methodology and workflow process for providing translational research informatics support. J Biomed Inform. 2009;42(2):377–81.

21. Institute CaLS. Perofrmance Standards for Antimicrobial Susceptiblity Testing. CLSI supplement M100. 2018.

22. Bolger AM, Lohse M, Usadel B. Trimmomatic: a flexible trimmer for Illumina sequence data. Bioinformatics. 2014;30(15):2114–20.

23. Bankevich A, Nurk S, Antipov D, Gurevich AA, Dvorkin M, Kulikov AS, et al. SPAdes: a new genome assembly algorithm and its applications to single-cell sequencing. J Comput Biol. 2012;19(5):455–77.

24. Kolmogorov M, Yuan J, Lin Y, Pevzner PA. Assembly of long, error-prone reads using repeat graphs. Nat Biotechnol. 2019;37(5):540–6.

25. Hunt M, Silva ND, Otto TD, Parkhill J, Keane JA, Harris SR. Circlator: automated circularization of genome assemblies using long sequencing reads. Genome Biol. 2015;16:294.

26. Vaser R, Sović, I., Nagarajan, N., & Šikić, M. Fast and accurate de novo genome assembly from long uncorrected reads. Genome research. 2017.

27. Page AJ, Cummins CA, Hunt M, Wong VK, Reuter S, Holden MT, et al. Roary: rapid large-scale prokaryote pan genome analysis. Bioinformatics. 2015;31(22):3691–3.

28. Löytynoja A. Phylogeny-aware alignment with PRANK. Totowa, NJ: Humana Press; 2014.

29. Stamatakis A. RAxML version 8: a tool for phylogenetic analysis and post-analysis of large phylogenies. Bioinformatics. 2014;30(9):1312–3.

30. Jolley KA, & Maiden, M. C. BIGSdb: scalable analysis of bacterial genome variation at the population level. BMC bioinformatics. 2010;11(1):595.

31. Seemann T. Prokka: rapid prokaryotic genome annotation. Bioinformatics. 2014;30(14):2068–9.

32. Jia B, Raphenya AR, Alcock B, Waglechner N, Guo P, Tsang KK, et al. CARD 2017: expansion and model-centric curation of the comprehensive antibiotic resistance database. Nucleic Acids Res. 2017;45(D1):D566–D73.

33. Carattoli A, Zankari E, Garcia-Fernandez A, Voldby Larsen M, Lund O, Villa L, et al. In silico detection and typing of plasmids using PlasmidFinder and plasmid multilocus sequence typing. Antimicrob Agents Chemother. 2014;58(7):3895–903.

34. Siguier P, Perochon J, Lestrade L, Mahillon J, Chandler M. ISfinder: the reference centre for bacterial insertion sequences. Nucleic Acids Res. 2006;34(Database issue):D32–6.

35. Hanson B, Johnson J, Leopold S, Sodergren E, Weinstock G. SVants – A long-read based method for structural variation detection in bacterial genomes. bioRxiv. 2019:822312.

36. Dumas JL, van Delden C, Perron K, Kohler T. Analysis of antibiotic resistance gene expression in Pseudomonas aeruginosa by quantitative real-time-PCR. FEMS Microbiol Lett. 2006;254(2):217–25.

37. Adler M, Anjum M, Berg OG, Andersson DI, Sandegren L. High fitness costs and instability of gene duplications reduce rates of evolution of new genes by duplication-divergence mechanisms. Mol Biol Evol. 2014;31(6):1526–35.

38. Gibson DG, Young L, Chuang RY, Venter JC, Hutchison CA, 3rd, Smith HO. Enzymatic assembly of DNA molecules up to several hundred kilobases. Nature methods. 2009;6(5):343–5.

39. Partridge SR, Zong Z, Iredell JR. Recombination in IS26 and Tn2 in the evolution of multiresistance regions carrying blaCTX-M-15 on conjugative IncF plasmids from Escherichia coli. Antimicrob Agents Chemother. 2011;55(11):4971–8.

40. Sandegren L, Linkevicius M, Lytsy B, Melhus A, Andersson DI. Transfer of an Escherichia coli ST131 multiresistance cassette has created a Klebsiella pneumoniae-specific plasmid associated with a major nosocomial outbreak. J Antimicrob Chemother. 2012;67(1):74–83.

41. Partridge SR. Analysis of antibiotic resistance regions in Gram-negative bacteria. FEMS Microbiol Rev. 2011;35(5):820–55.

42. Partridge SR, Kwong, S. M., Firth, N., S.O. Mobile Genetic Elements Associated with Antimicrobial Resistance. Clin Microbiol Rev. 2018;31(4):e00088–17.

43. He S, Hickman AB, Varani AM, Siguier P, Chandler M, Dekker JP, et al. Insertion Sequence IS26 Reorganizes Plasmids in Clinically Isolated Multidrug-Resistant Bacteria by Replicative Transposition. MBio. 2015;6(3):e00762.

44. Harmer CJ, Hall RM. IS26-Mediated Formation of Transposons Carrying Antibiotic Resistance Genes. mSphere. 2016;1(2).

45. Harmer CJ, Moran RA, Hall RM. Movement of IS26-associated antibiotic resistance genes occurs via a translocatable unit that includes a single IS26 and preferentially inserts adjacent to another IS26. MBio. 2014;5(5):e01801–14.

46. Anantham S, Hall RM. pCERC1, a small, globally disseminated plasmid carrying the dfrA14 cassette in the strA gene of the sul2-strA-strB gene cluster. Microb Drug Resist. 2012;18(4):364–71.

47. Boyd DA, Tyler S, Christianson S, McGeer A, Muller MP, Willey BM, et al. Complete nucleotide sequence of a 92-kilobase plasmid harboring the CTX-M-15 extended-spectrum beta-lactamase involved in an outbreak in long-term-care facilities in Toronto, Canada. Antimicrob Agents Chemother. 2004;48(10):3758–64.

48. Iida S, Mollet B, Meyer J, Arber W. Functional characterization of the prokaryotic mobile genetic element IS26. Molecular and General Genetics MGG. 1984;198(1):84–9.

49. Harmer CJ, Hall RM. IS26-Mediated Precise Excision of the IS26-aphA1a Translocatable Unit. MBio. 2015;6(6):e01866–15.

50. Livermore DM, Day M, Cleary P, Hopkins KL, Toleman MA, Wareham DW, et al. OXA-1 beta-lactamase and non-susceptibility to penicillin/beta-lactamase inhibitor combinations among ESBL-producing Escherichia coli. J Antimicrob Chemother. 2018.

51. Evans BA, Amyes SG. OXA beta-lactamases. Clin Microbiol Rev. 2014;27(2):241–63.

52. Evans SR, Hujer AM, Jiang H, Hujer KM, Hall T, Marzan C, et al. Rapid Molecular Diagnostics, Antibiotic Treatment Decisions, and Developing Approaches to Inform Empiric Therapy: PRIMERS I and II. Clin Infect Dis. 2016;62(2):181–9.

53. Bevan ER, Jones AM, Hawkey PM. Global epidemiology of CTX-M beta-lactamases: temporal and geographical shifts in genotype. J Antimicrob Chemother. 2017;72(8):2145–55.

54. Mena A, Plasencia V, Garcia L, Hidalgo O, Ayestaran JI, Alberti S, et al. Characterization of a large outbreak by CTX-M-1-producing Klebsiella pneumoniae and mechanisms leading to in vivo carbapenem resistance development. J Clin Microbiol. 2006;44(8):2831–7.

55. Stoesser N, Sheppard AE, Pankhurst L, De Maio N, Moore CE, Sebra R, et al. Evolutionary History of the Global Emergence of the Escherichia coli Epidemic Clone ST131. mBio. 2016;7(2):e02162.

56. Poirel L, Lartigue M-F, Decousser J-W, Nordmann P. ISEcp1B-mediated transposition of blaCTX-M in Escherichia coli. Antimicrobial agents and chemotherapy. 2005;49(1):447–50.

57. Zowawi HM, Forde BM, Alfaresi M, Alzarouni A, Farahat Y, Chong TM, et al. Stepwise evolution of pandrug-resistance in Klebsiella pneumoniae. Sci Rep. 2015;5:15082.

58. Simner PJ, Antar AAR, Hao S, Gurtowski J, Tamma PD, Rock C, et al. Antibiotic pressure on the acquisition and loss of antibiotic resistance genes in Klebsiella pneumoniae. J Antimicrob Chemother. 2018;73(7):1796–803.

59. Hernández-Allés S, Benedí VJ, Martínez-Martínez L, Pascual Á, Aguilar A, Tomás JM, et al. Development of resistance during antimicrobial therapy caused by insertion sequence interruption of porin genes. Antimicrobial agents and chemotherapy. 1999;43(4):937–9.

60. Doumith M, Ellington MJ, Livermore DM, Woodford N. Molecular mechanisms disrupting porin expression in ertapenem-resistant Klebsiella and Enterobacter spp. clinical isolates from the UK. J Antimicrob Chemother. 2009;63(4):659–67.

61. Vandecraen J, Chandler M, Aertsen A, Van Houdt R. The impact of insertion sequences on bacterial genome plasticity and adaptability. Crit Rev Microbiol. 2017;43(6):709–30.

62. Sandegren L, Andersson DI. Bacterial gene amplification: implications for the evolution of antibiotic resistance. Nat Rev Microbiol. 2009;7(8):578–88.

63. Andersson DI, Hughes D. Antibiotic resistance and its cost: is it possible to reverse resistance? Nat Rev Microbiol. 2010;8(4):260–71.

64. Leavitt A, Chmelnitsky I, Colodner R, Ofek I, Carmeli Y, Navon-Venezia S. Ertapenem resistance among extended-spectrum-beta-lactamase-producing Klebsiella pneumoniae isolates. J Clin Microbiol. 2009;47(4):969–74.

65. Nicoloff H, Hjort K, Levin BR, Andersson DI. The high prevalence of antibiotic heteroresistance in pathogenic bacteria is mainly caused by gene amplification. Nat Microbiol. 2019;4(3):504–14.

66. Gullberg E, Albrecht LM, Karlsson C, Sandegren L, Andersson DI. Selection of a multidrug resistance plasmid by sublethal levels of antibiotics and heavy metals. mBio. 2014;5(5):e01918–14.

67. Andersson DI, Nicoloff H, Hjort K. Mechanisms and clinical relevance of bacterial heteroresistance. Nat Rev Microbiol. 2019;17(8):479–96.

