## Supplemental Material for "Concurrence of Porin Loss and Modular Amplification of β-Lactamase Encoding Genes Drives Carbapenem Resistance in a Cohort of Recurrent *Enterobacterales* Bacteremia"

**LIST OF SUPPLEMENTAL MATERIALS:**

Supplementary Tables

Table S1. Oxford Nanopore Technology + Illumina Assembly QC Results

Table S2: Primers and Probes for qRT-PCR and Cloning

Table S3. Clinical and Demographic Characteristics of Patients with and without Recurrent ESBL-E Bacteremia

Table S4. Summary of sequence types and β-lactamase encoding genes present in studied isolates

Table S5. ST131 Serial Isolates Coverage Depth Analysis

Table S6. Patient 10 Serial Isolates Coverage Depth Analysis

Supplementary Figures

FIG S1. Assessment of clonality among serial isolates.

FIG S2. Genomic context of β-lactamase encoding genes harbored on translocatable units in patient 11 strains p11A and p11B.

Fig S3. Alignment of Tn*MB1860* chromosomal context with ST131 isolates with p11A multireplicon F type plasmid locus containing *bla*$_{\text{OXA-1}}$ and *bla*$_{\text{CTX-M-15}}$.

FIG S4. *Bla*$_{\text{OXA-1}}$ expression contributes to piperacillin-tazobactam (TZP) resistance.

Fig S5. Serial passage of p4C (A) and p4D (B) strains in absence of ETP selective pressure over 10-day period.

### Table S1. Oxford Nanopore Technology + Illumina Assembly QC Results

| MDACC ID | Isolate | Species | ST | Assembly | ONT Average Coverage | # of contigs | Total bp | Average GC% | BioSample Accession# |
|---|---|---|---|---|---|---|---|---|---|
| MB101 | p101 | *K. pneumoniae* | 37 | Incomplete | 193 | 8 | 5591454 | 57.0 | SAMN07189586 |
| MB746A | p746 | *E. coli* | 405 | Incomplete | 218 | 6 | 5525438 | 50.5 | SAMN07189563 |
| MB1860 | p4A | *E. coli* | 131 | Complete | 206 | 1 | 5255498 | 50.7 | SAMN13948677 |
| MB2374 | p4B | *E. coli* | 131 | Incomplete | 227 | 2 | 5315955 | 50.7 | SAMN13948678 |
| MB2463 | p4C | *E. coli* | 131 | Incomplete | 199 | 2 | 5329559 | 50.7 | SAMN13948679 |
| MB2573 | p4D | *E. coli* | 131 | Incomplete | 163 | 2 | 5345805 | 50.7 | SAMN13948680 |
| MB2315 | p10A | *E. coli* | 10 | Complete | 384 | 5 | 4920692 | 50.8 | SAMN13948681 |
| MB2446 | p10B | *E. coli* | 10 | Complete | 263 | 5 | 4905205 | 50.8 | SAMN13948682 |
| MB2649 | p10C | *E. coli* | 10 | Complete | 428 | 5 | 4879573 | 50.8 | SAMN13948683 |
| EC215 | p11A | *E. coli* | 131 | Complete | 511 | 3 | 5545837 | 50.7 | SAMN13948684 |
| MB2489 | p11B | *E. coli* | 131 | Incomplete | 71 | 3 | 5580474 | 50.7 | SAMN13948685 |
| MB1860_1 | p4A_1 | *E. coli* | 131 | Complete | 410 | 1 | 5280799 | 50.7 | SAMN13948686 |
| MB1860_2 | p4A_2 | *E. coli* | 131 | Complete | 501 | 1 | 5304001 | 50.7 | SAMN13948687 |
| MB1860_3 | p4A_3 | *E. coli* | 131 | Complete | 176 | 3 | 5279381 | 50.7 | SAMN13948688 |
| MB1860_4 | p4A_4 | *E. coli* | 131 | Complete | 381 | 2 | 5328184 | 50.7 | SAMN13948689 |
| MB1860_H1 | p4A_H1 | *E. coli* | 131 | Complete | 150 | 1 | 5296385 | 50.7 | SAMN13948690 |
| MB1860_H2 | p4A_H2 | *E. coli* | 131 | Complete | 311 | 1 | 5324283 | 50.7 | SAMN13948691 |

### Table S2: Primers and Probes for qRT-PCR and Cloning

| Gene | Foward Primer | Reverse Primer | Probe |
|---|---|---|---|
| *bla*$_{\text{CTX-M}}$ for qRT-PCR | 5’ – ACCGAGCCGACGTTAAACAC – 3’ | 5’ – TGCCCGAGGTGAAGTGGTAT – 3’ | CATTCCGGGCGATCCGCG |
| *bla*$_{\text{OXA-1}}$ for qRT-PCR | 5’ – AAAACCCCCAAAGGAATGG – 3’ | 5’ – AAACCCAAACAACAGAAAATTG – 3’ | CTGGAACAGCAATCATACACCAAAGACGTG |
| *rpsL* for qRT-PCR | 5’ – AACCACGTGCTCGCAAAGTT – 3’ | 5’ – GTTTTTGCGGGCATGCTT – 3’ | CGAAAAGCAACGTGCCTGCGC |
| *bla*$_{\text{CTX-M-15}}$ for cloning | 5’ – GTACCCGGGGATCCTCTAGACCAGAATAAGGAATCCCATGG – 3’ | 5’ – GCTTGCATGCCTGCAGTTTCCCCATTCCGTTTCCG – 3’ | |
| *bla*$_{\text{OXA-1}}$ for cloning | 5’ – GTACCCGGGGATCCTCTAGACCGTTAAAATTAAGCCCTTTACC – 3’ | 5’ – GCTTGCATGCCTGCAGAAGGGTTGGGCGATTTTGC – 3’ | |

**Table S3. Clinical and Demographic Characteristics of Patients with and without Recurrent ESBL-E Bacteremia**

| Characteristics | No recurrence (n = 100)[a] | Recurrence (n = 16)[a] | p-value[a] |
|---|---|---|---|
| Age (years)[b] | 60 (45 - 66) | 50 (29 - 66) | 0.36 |
| Gender (male) | 58 (59) | 8 (50) | 0.59 |
| Treatment service | | | 0.07 |
| *Leukemia* | 37 (37) | 12 (75) | |
| *Stem cell transplant* | 19 (19) | 2 (13) | |
| *Lymphoma/myeloma* | 11 (11) | 0 (0) | |
| *GI medical oncology* | 15 (15) | 0 (0) | |
| *Other* | 18 (18) | 2 (13) | |
| ANC (cells/µl)[b,c] | 790 (0 - 8850) | 0 (0 - 150) | < 0.01 |
| Organism | | | 1.00 |
| *E. coli* | 86 (86) | 14 (88) | |
| *K. pneumoniae* | 12 (12) | 2 (12) | |
| *K. oxytoca* | 2 (2) | 0 (0) | |

[a]All values n (%) are compared using Fisher's exact test unless otherwise listed.

[b]Median, interquartile range (tested with Wilcoxon rank-sum test)

[c]ANC: absolute neutrophil count, normal is >1,500 cells/µl

| Table S4. Summary of sequence types and β-lactamase encoding genes present in studied isolates | | | | | | | | | | | |
|---|---|---|---|---|---|---|---|---|---|---|---|
| **Species** | **Strain** | **Patient #** | **Isolate #** | **ST** | **β-lactamase encoding genes** | **CAZ MIC* (S, ≤ 4 I, 8 R, ≥ 16)** | **CEP MIC* (S, ≤ 2 I, 4-8 R, ≥ 16)** | **TZP MIC* (S, ≤ 16 I, 32-64 R, ≥ 128)** | **ETP MIC* (S, ≤ 0.5 I, 4-8 R, ≥ 16)** | **MEM MIC* (S, ≤ 1 I, 2 R, ≥ 4)** | **Increased mapping density of β-lactamase-encoding genes%** |
| *E. coli* | MB1159 | 1 | A | 131 | $bla_{OXA-1}$, $bla_{CTX-M-15}$ | 4 | 2 | ≤ 4 | ≤0.5 | ≤0.25 | None |
| | MB1283 | | B | 131 | $bla_{OXA-1}$, $bla_{CTX-M-15}$ | 4 | 2 | ≤ 4 | ≤0.5 | ≤0.25 | None |
| | MB1860 | 4 | A | 131 | $bla_{OXA-1}$, $bla_{CTX-M-15}$ | 16 | ≥64 | 8 | ≤0.5 | ≤0.25 | None |
| | MB2374 | | B | 131 | $bla_{OXA-1}$, $bla_{CTX-M-15}$ | 16 | 8 | ≥128 | ≤0.5 | ≤0.25 | $bla_{OXA-1}$ |
| | MB2463 | | C | 131 | $bla_{OXA-1}$, $bla_{CTX-M-15}$ | 16 | 64 | ≥128 | ≥32 | 4 | $bla_{OXA-1}$ |
| | MB2573 | | D | 131 | $bla_{OXA-1}$, $bla_{CTX-M-15}$ | ≥64 | ≥64 | ≥128 | ≥32 | 4 | $bla_{OXA-1}$ |
| | MB1341 | 5 | A | 131 | $bla_{TEM-1}$, $bla_{CTX-M-14}$ | 4 | ≥64 | 64 | ≤0.5 | ≤0.25 | $bla_{TEM-1}$ |
| | MB1676 | | B | 1604 | None | ≤ 1 | ≤ 1 | ≤4 | ≤0.5 | ≤0.25 | None |
| | MB1787 | 6 | A | 156 | $bla_{TEM-169}$, $bla_{CTX-M-15}$ | ≥ 64 | ≥ 64 | ≥128 | ≤0.5 | ≤0.25 | $bla_{TEM-169}$, $bla_{CTX-M-15}$ |

| | | | | | | | | | | | |
|---|---|---|---|---|---|---|---|---|---|---|---|
| | MB1871B | | B | 156 | *bla*$_{\text{TEM-169}}$, *bla*$_{\text{CTX-M-15}}$ | ≥ 64 | ≥ 64 | ≥128 | ≤0.5 | ≤0.25 | *bla*$_{\text{TEM-169}}$, *bla*$_{\text{CTX-M-15}}$ |
| | MB1287 | 7 | A | 405 | *bla*$_{\text{SHV-12}}$, *bla*$_{\text{CTX-M-15}}$ | ≥64 | 16 | ≤ 4 | ≤0.5 | ≤ 0.25 | None |
| | MB1257 | | B | 405 | *bla*$_{\text{OXA-1}}$, *bla*$_{\text{CTX-M-15}}$ | 16 | 16 | 16 | ≤ 0.5 | ≤ 0.25 | None |
| | MB1257A | | C | 10 | *bla*$_{\text{OXA-181}}$ | ≥ 64 | 16 | ≥128 | ≥ 32 | ≥ 16 | None |
| | MB1339 | 8 | A | 10 | *bla*$_{\text{OXA-1}}$, *bla*$_{\text{CTX-M-15}}$ | ≥ 64 | ≥ 64 | ≥128 | ≤ 0.5 | ≤ 0.25 | *bla*$_{\text{OXA-1}}$, *bla*$_{\text{CTX-M-15}}$ |
| | MB1418 | | B | 2659 | *bla*$_{\text{CTX-M-55}}$ | 4 | 2 | ≤4 | ≤ 0.5 | ≤ 0.25 | None |
| | MB1248 | 9 | A | 744 (ST10 like) | *bla*$_{\text{CTX-M-55}}$ | 16 | 2 | ≤4 | ≤ 0.5 | ≤ 0.25 | None |
| | MB1562 | | B | 744 (ST10 like) | *bla*$_{\text{CTX-M-55}}$ | 16 | 2 | ≤4 | ≤ 0.5 | ≤ 0.25 | None |
| | MB2315 | 10 | A | 10 | *bla*$_{\text{CTX-M-55}}$ | 16 | 2 | 8 | ≤ 0.5 | ≤ 0.25 | No |
| | MB2446 | | B | 10 | *bla*$_{\text{CTX-M-55}}$ | ≥ 64 | ≥ 64 | ≥128 | ≥ 32 | 8 | *bla*$_{\text{CTX-M-55}}$ |
| | MB2649 | | C | 10 | *bla*$_{\text{CTX-M-55}}$ | ≥ 64 | ≥ 64 | 16 | ≤ 0.5 | ≤ 0.25 | *bla*$_{\text{CTX-M-55}}$ |
| | EC215 | 11 | A | 131 | *bla*$_{\text{OXA-1}}$, *bla*$_{\text{CTX-M-15}}$ | ≥ 64 | ≥ 64 | 64 | ≤ 0.5 | ≤ 0.5 | *bla*$_{\text{OXA-1}}$, *bla*$_{\text{CTX-M-15}}$ |
| | MB2489 | | B | 131 | *bla*$_{\text{OXA-1}}$, *bla*$_{\text{CTX-M-15}}$ | ≥ 64 | ≥ 64 | ≥128 | 4 | 1 | *bla*$_{\text{OXA-1}}$, *bla*$_{\text{CTX-M-15}}$ |
| | MB746A | N/A | N/A | 405 | *bla*$_{\text{OXA-1}}$, *bla*$_{\text{CTX-M-15}}$ | 64 | ≥64 | ≥128 | ≥32 | 4 | *bla*$_{\text{OXA-1}}$ |

| | | | | | | | | | | | |
|---|---|---|---|---|---|---|---|---|---|---|---|
| *K. pneumoniae* | KP3076 | 2 | A | Novel | *bla*$_{\text{SHV-1}}$, *bla*$_{\text{TEM-1}}$, *bla*$_{\text{OXA-1}}$, *bla*$_{\text{CTX-M-15}}$ | 4 | 2 | 16 | ≤ 0.5 | ≤ 0.25 | None |
| | MB1927^ | | B | Novel | *bla*$_{\text{LEN}}$ | ≤ 1 | ≤ 1 | ≤4 | ≤ 0.5 | ≤ 0.25 | None |
| | MB1647 | 3 | A | 25 | *bla*$_{\text{SHV-11}}$, *bla*$_{\text{TEM-1}}$, *bla*$_{\text{OXA-1}}$, *bla*$_{\text{CTX-M-15}}$ | 16 | 8 | ≥128 | ≤ 0.5 | ≤ 0.25 | None |
| | MB3087 | | B | 25 | *bla*$_{\text{SHV-11}}$, *bla*$_{\text{TEM-1}}$, *bla*$_{\text{OXA-1}}$, *bla*$_{\text{CTX-M-15}}$ | 16 | 8 | ≥128 | ≤ 0.5 | ≤ 0.25 | None |
| | MB101 | N/A | N/A | 37 | *bla*$_{\text{SHV-11}}$, *bla*$_{\text{TEM-1}}$, *bla*$_{\text{OXA-1}}$, *bla*$_{\text{CTX-M-15}}$ | ≥64 | ≥64 | ≥128 | ≥32 | 8 | *bla*$_{\text{OXA-1}}$, *bla*$_{\text{CTX-M-15}}$ |

Abbreviations: ST = sequence type, MIC = minimum inhibitory concentration, CAZ = ceftazidime, CEP = cefepime, TZP = piperacillin-tazobactam, ERT = ertapenem, MEM = meropenem, *MIC reported in µg/mL with S, I, R referring to susceptible, intermediate, resistant as considered by CLSI, ^reported as *K. pneumoniae* by clinical laboratory but most closely matches *K. varicola* by WGS. %Increased mapping density defined as > 2-fold relative to multi-locus sequence typing genes.

### Table S5. ST131 Serial Isolates Coverage Depth Analysis

| | | Sample ID | Average Fold Coverage[a] | | | | Estimated Gene Copy Number | | |
|---|---|---|---|---|---|---|---|---|---|
| | | | MLST | $bla_{CTX-M-15}$ | $bla_{OXA-1}$ | *repB* FIA | $bla_{CTX-M-15}$ | $bla_{OXA-1}$ | *repB* FIA |
| Patient 4 | Illumina | p4A | 352.87 | 320.73 | 379.47 | NA | 0.91 | 1.08 | NA |
| | | p4B | 135.80 | 132.38 | 1584.12 | NA | 0.97 | 11.66 | NA |
| | | p4C | 84.95 | 74.39 | 1557.87 | NA | 0.88 | 18.34 | NA |
| | | p4D | 217.07 | 258.51 | 5947.84 | NA | 1.19 | 27.40 | NA |
| | ONT | p4A | 205.43 | 156.75 | 182.64 | NA | 0.76 | 0.89 | NA |
| | | p4B | 223.86 | 159.74 | 1548.03 | NA | 0.71 | 6.92 | NA |
| | | p4C | 196.04 | 146.69 | 1519.44 | NA | 0.75 | 7.75 | NA |
| | | p4D | 164.60 | 125.59 | 1959.17 | NA | 0.76 | 11.90 | NA |
| Patient 11 | Illumina | p11A | 199.98 | 442.39 | 592.56 | 280.68 | 2.21 | 2.96 | 1.40 |
| | | p11B | 85.71 | 208.39 | 315.43 | 139.64 | 2.43 | 3.68 | 1.63 |
| | ONT | p11A | 485.86 | 850.98 | 884.72 | 392.94 | 1.75 | 1.82 | 0.81 |
| | | p11B | 67.48 | 98.58 | 109.01 | 68.16 | 1.46 | 1.62 | 1.01 |

[a]See Materials and Methods for Calculation

### Table S6. Patient 10 Serial Isolates Coverage Depth Analysis

| | Sample ID | Average Fold Coverage[a] | | | | | Estimated Gene Copy Number | | | |
|---|---|---|---|---|---|---|---|---|---|---|
| | | MLST | $bla_{CTX-M-55}$ | RNA | *sul2* | *repB* FIC | $bla_{CTX-M-55}$ | RNA | *sul2* | *repB* FIC |
| Illumina | p10A | 55.45 | 78.56 | 471.05 | 269.93 | 89.12 | 1.42 | 8.49 | 4.87 | 1.61 |
| | p10B | 168.42 | 173.92 | 1836.21 | 38.63 | 65.90 | 1.03 | 10.90 | 0.23 | 0.39 |
| | p10C | 228.86 | 1513.11 | 1669.76 | 1052.54 | 334.28 | 6.61 | 7.30 | 4.60 | 1.46 |
| ONT | p10A | 373.49 | 208.81 | 466.84 | 298.52 | 180.65 | 0.56 | 1.25 | 0.80 | 0.48 |
| | p10B | 257.71 | 1182.41 | 2818.97 | 331.18 | 114.72 | 4.59 | 10.94 | 1.29 | 0.45 |
| | p10C | 419.54 | 9476.34 | 17888.10 | 7757.20 | 168.70 | 22.59 | 42.64 | 18.49 | 0.40 |

[a]See Materials and Methods for Calculation

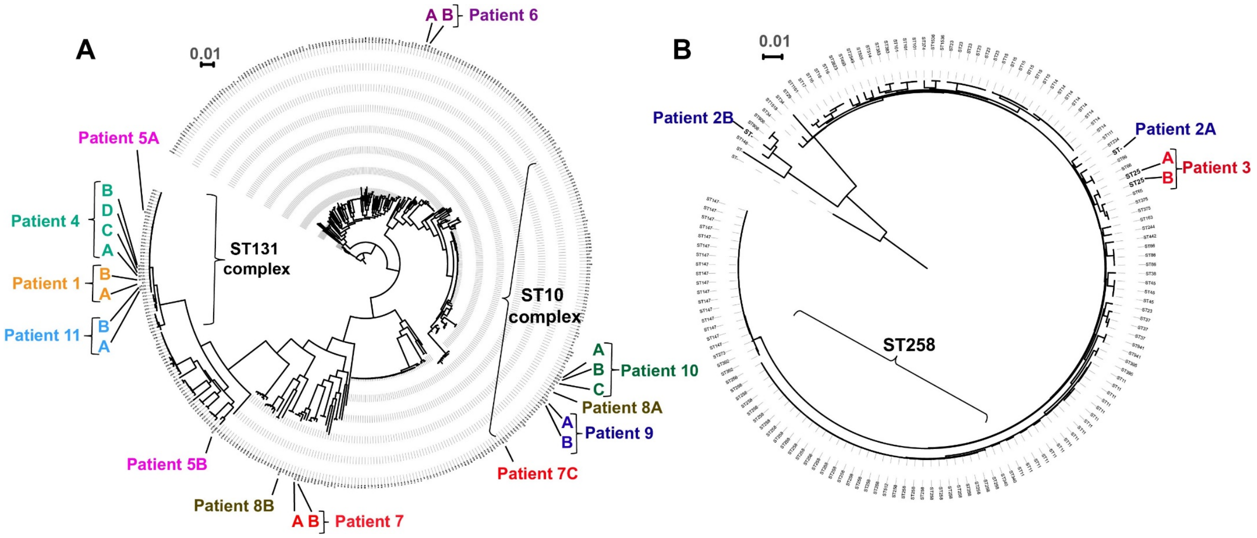


**FIG S1** Assessment of clonality among serial isolates. Scalebar indicates 0.01 nucleotide substitutions per site. Major STs are labeled for reference purposes. **(A)** Maximum likelihood phylogenetic tree of publicly available *E. coli* strains combined with strains from this cohort. Patient isolates are color coded and labeled. When strains from a particular patient were co-localized (e.g. patient 6), they were considered likely clonal. When strains were on distinct parts of the phylogenetic tree (e.g. patient 5), they were considered likely non-clonal. **(B)** Phylogenetic tree and clonal analysis as described in (A) for *K. pneumoniae* isolates.

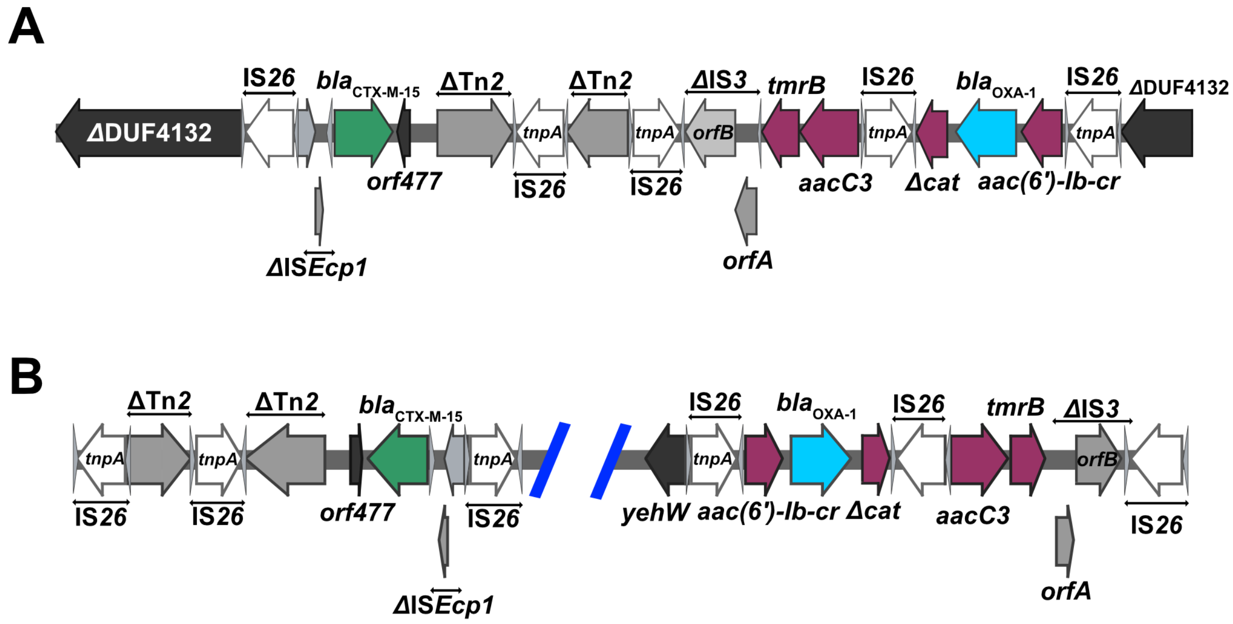


**FIG S2** Genomic context of β-lactamase encoding genes harbored on translocatable units in patient 11 strains p11A and p11B. **(A)** The chromosomal context of *TnMB1860* (GenBank Accession #: CP049077) on p11 and p11B. Open reading frames (ORFs) and their transcriptional orientation are represented by arrows with respective annotations. Terminal left and right inverted repeats ($IR_L$ and $IR_R$ respectively) of ISs are specified by grey triangles that bracket respective complete and incomplete *tnpA* genes. ORFs are colored as follows: AMR genes (maroon), $bla_{OXA-1}$ (blue), $bla_{CTX-M-15}$ (green), IS*26 tnpA* (white), and other IS/Tn elements (gray). Delta (Δ) next to annotated genetic region indicates a truncation or disruption**. (B)** Genomic context of the $bla_{OXA-1}$ (blue) and $bla_{CTX-M-15}$ (green) genes on a multireplicon, ~180 kbp, F type plasmid (GenBank Accession #: CP049079) carried by p11A and p11B.

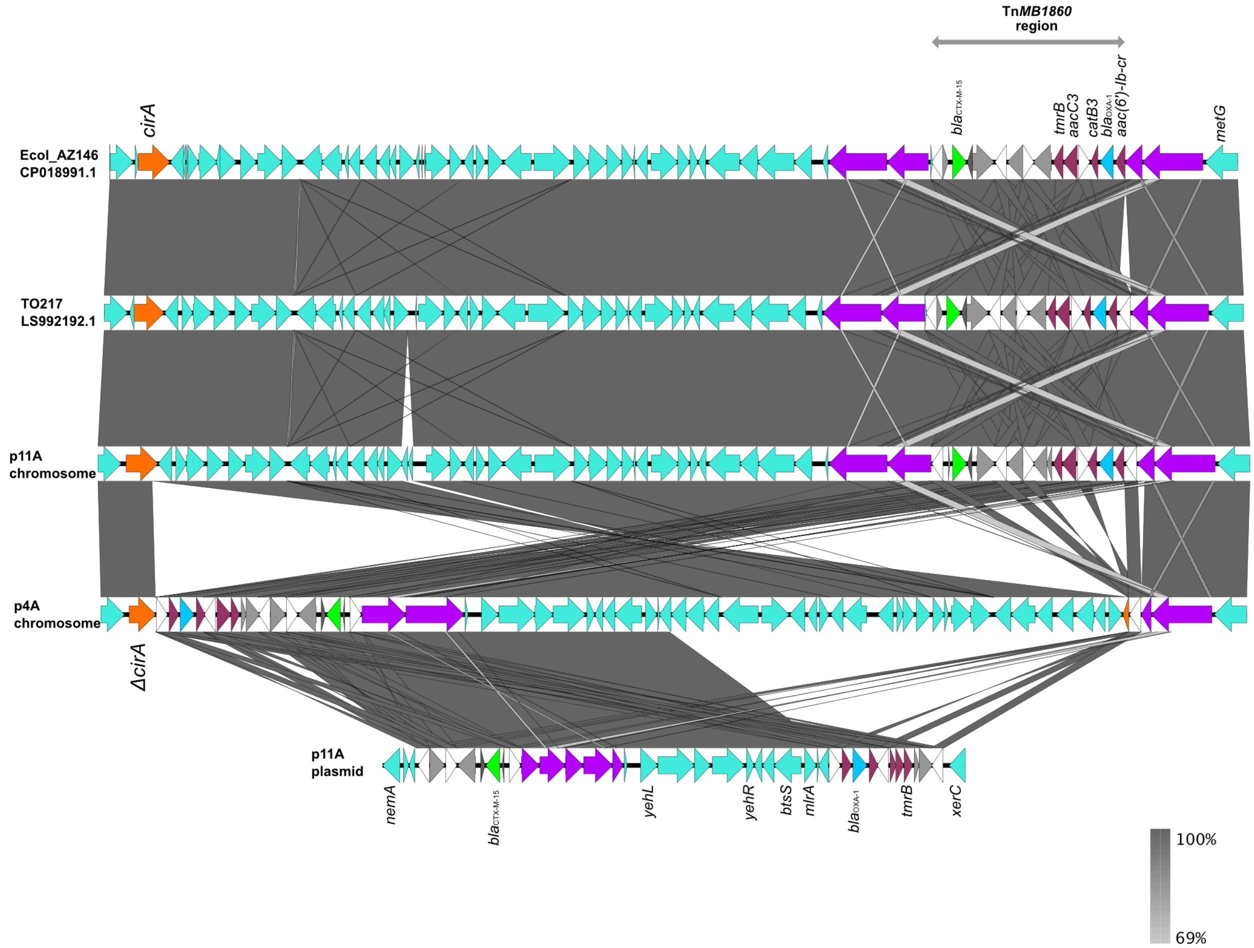


**Fig S3** Alignment of Tn*MB1860* chromosomal context with ST131 isolates with p11A multireplicon F type plasmid locus containing *bla*$_{\text{OXA-1}}$ and *bla*$_{\text{CTX-M-15}}$. *bla*$_{\text{OXA-1}}$ ORF is in blue, *bla*$_{\text{CTX-M-15}}$ is in green, IS26 *tnpA* is in white, Tn2-like ORFs in grey, *cirA* in orange, DUF4132 regions in purple, and other ORFs in turquoise. The Tn*MB1860* region in Ecol_AZ146 is noted by the grey, double-headed arrow at the top. GenBank Accession numbers are CP0018991.1 and LS992192.1 for Ecol_AZ146 and TO217 respectively. GenBank Accession numbers for p4A chromosome, p11A chromosome, and p11A plasmid are CP049085, CP049077, and CP049079 respectively.

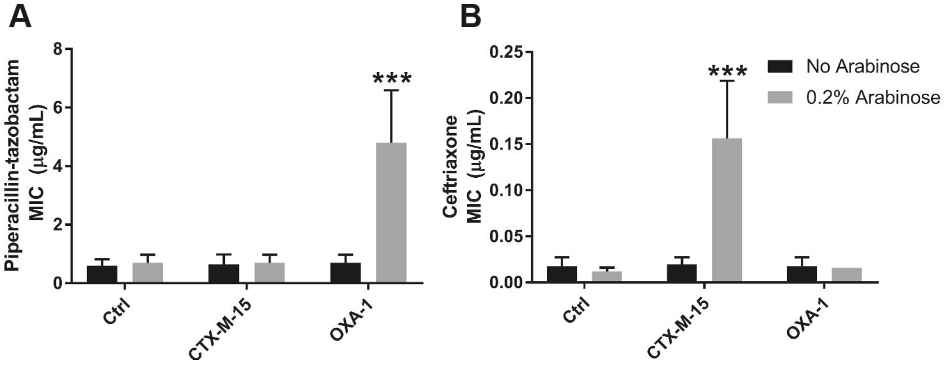


**FIG S4** *Bla*$_{\text{OXA-1}}$ expression contributes to piperacillin-tazobactam (TZP) resistance. DH5α *E. coli* carrying pBAD33 empty vector (Ctrl) or arabinose-inducible *bla*$_{\text{OXA-1}}$ or *bla*$_{\text{CTX-M-15}}$ were exposed to, piperacillin-tazobactam **(A)** or ceftriaxone **(B)** in a broth microdilution assay. Mean and standard deviation of MICs observed from at least three experiments are reported. Statistics reported as results of ANOVA with Dunnett's test of multiple comparisons against uninduced controls. *P*-value of $<0.0001$ shown as ***

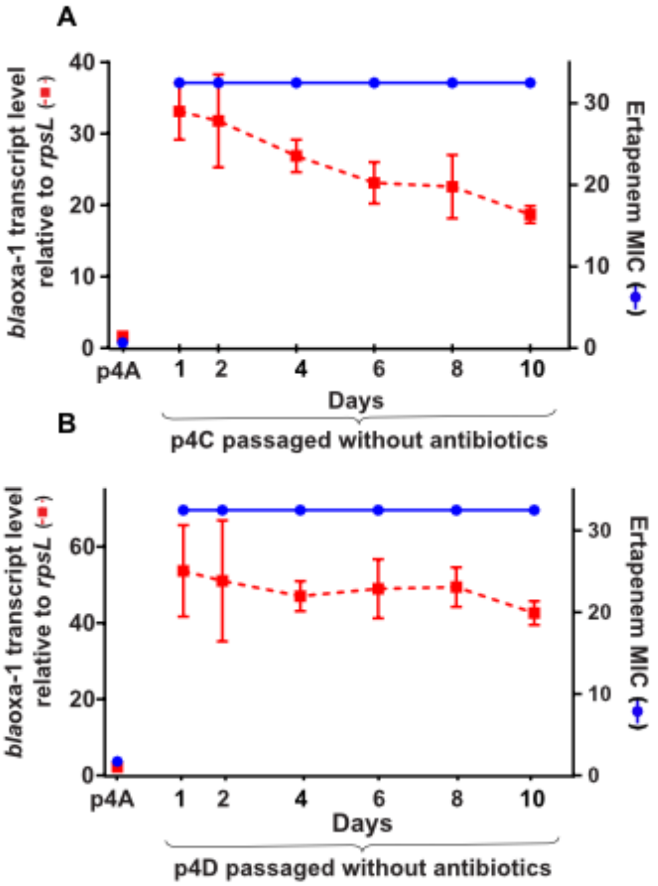


**Fig S5** Serial passage of p4C **(A)** and p4D **(B)** strains in absence of ETP selective pressure over 10-day period. Left axis indicates relative copy number of $bla_{\mathrm{OXA\text{-}1}}$ shown in red while right axis indicates ETP MIC (μg/ml) in blue. Parental strain p4D results shown in bottom left corner.
